# Distinct neocortical mechanisms underlie human SI responses to median nerve and laser evoked peripheral activation

**DOI:** 10.1101/2021.10.11.463545

**Authors:** Ryan V. Thorpe, Christopher J. Black, David A. Borton, Li Hu, Carl Y. Saab, Stephanie R. Jones

**Affiliations:** Department of Neuroscience, Brown University, Providence, RI, United States; School of Engineering, Brown University, Providence, RI, United States; Center for Neurorestoration and Neurotechnology, Providence VAMC, Providence, RI, United States; Key Laboratory of Mental Health, Institute of Psychology, Chinese Academy of Sciences, Beijing 100101, China; Department of Neurosurgery, Rhode Island Hospital, Providence, RI, United States

## Abstract

Magneto- and/or electro-encephalography (M/EEG) are non-invasive clinically-relevant tools that have long been used to measure electromagnetic fields in somatosensory cortex evoked by innocuous and noxious somatosensory stimuli. Two commonly applied stimulation paradigms that produce distinct responses in primary somatosensory cortex (SI) linked to innocuous and noxious sensations are electrical median nerve (MN) stimulation and cutaneous laser-evoked (LE) stimulation to the dorsum of the hand, respectively. Despite their prevalence, the physiological mechanisms that produce stereotypic macroscale MN and LE responses have yet to be fully articulated, limiting their utility in understanding brain dynamics associated with non-painful and/or painful somatosensation. Through a literature review, we detailed features of MN and LE responses source-localized to SI that are robust and reproducible across studies. We showed that the first peak in the MN response at ∼20 ms post-stimulus (i.e., MN N1) corresponds to outward-directed deep-to-superficial electrical current flow through the cortical laminae, which is followed by inward-directed current at ∼30 ms (i.e., MN P1). In contrast, the initial LE response occurs later at ∼170 ms (i.e., LE N1) and is oriented inward and opposite the direction of the MN N1. We then examined the neocortical circuit mechanisms contributing to the robust features of each response using the Human Neocortical Neurosolver (HNN) neural modeling software tool (Neymotin et al., 2020). Using HNN as a hypothesis development and testing tool, model results predicted the MN response can be simulated with a sequence of layer specific thalamocortical and cortico-cortical synaptic drive similar to that previously reported for tactile evoked responses (Jones et al., 2007; Neymotin et al., 2020), with the novel discovery that an early excitatory input to supragranular layers at ∼30 ms is an essential mechanism contributing to the inward current flow of the MN P1. Model results further predicted that the initial ∼170 ms inward current flow of the LE N1 was generated by a burst of repetitive gamma-frequency (∼40 Hz) excitatory synaptic drive to supragranular layers, consistent with prior reports of LE gamma-frequency activity. These results make novel and detailed multiscale predictions about the dynamic laminar circuit mechanisms underlying temporal and spectral features of MN and LE responses in SI and can guide further investigations in follow-up studies. Ultimately, these findings may help with the development of targeted therapeutics for pathological somatosensation, such as somatic sensitivity and acute neuropathic pain.

## 1. Introduction

Somatosensation is a crucial mechanism for healthy human behavior and survival that is often segregated into two categories: non-painful (e.g., gentle touch or peripheral nerve stimulation) and painful (e.g., unpleasant heat) sensations. The distinct neural receptors and pathways mediating innocuous and noxious somatosensory signals to the brain have been well categorized (Almeida et al., 2004; Burke et al., 1975; Lamour et al., 1983; Treede et al., 1999; Willis, 2007, 1985). However, how these signals are uniquely processed within brain circuits to generate reciprocal sensations of innocuous, non-nociceptive somatosensation *versus* noxious, nociceptive somatosensation is not well understood. The brain’s electrophysiological correlates of somatosensation can be readily studied non-invasively in humans using electro- and/or magneto-encephalography (M/EEG), which reflects fast-time scale neural signals from neocortex. Numerous studies have identified unique and robust M/EEG-measured evoked responses to non-painful and painful somatosensory stimuli, yet, little is known of the dynamic, multiscale neocortical circuits that produce these perceptually distinct evoked responses.

Two respective innocuous and noxious peripheral stimulation paradigms stand out in the literature as producing distinct responses in primary somatosensory cortex (SI): electrical median nerve (MN) stimulation and cutaneous laser-evoked (LE) stimulation to the dorsum of the hand. Although MN and laser stimuli are not naturalistic sensory stimuli, uncovering the underlying neural processes they induce can provide a foundation for understanding how the brain differentially processes non-painful *versus* painful sensations in general (Lenoir et al., 2017). MN stimulation, and more broadly tactile stimulation of the hand, evokes its earliest electrophysiological response in contralateral SI, specifically area 3b/1, at ∼20 ms (Allison et al., 1989; Cauller and Kulics, 1991; Huttunen et al., 2006; Inui et al., 2004; Kakigi et al., 2000; Kanno et al., 2003; Lin et al., 2005, 2003; Lipton et al., 2006; Schroeder et al., 1995; Tiihonen et al., 1989; Wikström et al., 1996). Laser stimulation of the hand evokes a later response in a topographically similar region of contralateral SI at ∼170 ms (Christmann et al., 2007; Jin et al., 2018; Kakigi et al., 2005; Ploner et al., 1999; Timmermann et al., 2001; Valentini et al., 2012). While different sub-second time-domain components (e.g., peak amplitudes) of the MN response and LE response in SI appear to aid in encoding the location (Cauller and Kulics, 1991; Ishibashi et al., 2000; Omori et al., 2013; Schroeder et al., 1995), strength, and perceived intensity (Lin et al., 2003; Timmermann et al., 2001) of the sensory stimulus, the precise neural mechanisms that facilitate such an encoding remain unknown.

Here, we first identify distinct components of the source-localized MN response (Forss et al., 2001; Inui et al., 2004; Kanno et al., 2003; Lin et al., 2005, 2003, 2003; Sorrentino et al., 2009; Tiihonen et al., 1989; Wikström et al., 1996; Zainea et al., 2005) compared with the source-localized LE response (Ahmed Mahmutoglu et al., 2022; Baumgärtner et al., 2011; Jin et al., 2018; Omori et al., 2013; Ploner et al., 1999; Raij et al., 2003; Schlereth et al., 2003; Valentini et al., 2012), as reported widely across prior studies. Given these distinctions and the aforementioned observation that the MN and LE responses both originate from a similar region in SI, we use a biophysically constrained neural circuit model via the Human Neocortical Neurosolver (HNN) modeling platform to test a series of targeted hypotheses on *how* the MN and LE responses are generated in SI through different cell and circuit mechanisms of a common underlying neocortical circuit. In brief, HNN is a user-friendly software tool that simulates the primary electrical currents responsible for generating source-localized M/EEG signals in a biophysically-principled laminar neocortex model (Neymotin et al., 2020). HNN is designed to develop targeted and testable predictions on the multiscale neural mechanisms of current dipole sources from a localized region of cortex. HNN has been used to study circuit mechanisms of innocuous tactile stimulation to the first digit of the hand and correlates of tactile perception in SI measured with MEG and EEG (Jones et al., 2009, 2007; Law et al., 2021; Neymotin et al., 2020; Sliva et al., 2018; Ziegler et al., 2010). We begin the modeling portion of this study by first extending the simulated cell and circuit mechanisms of the tactile evoked (TE) response to the MN response, which have common afferent pathways and similar evoked response profiles. We then expand our modeling investigation to the LE response.

By testing and refining mechanistic hypotheses based on prior literature while optimizing the output of the model to recorded current dipole source data, we show that waveform features of the respective MN and LE responses emerge from distinct cell and circuit generation mechanisms. Several of these predictions were further supported by studies not used to constrain the model investigation, and other predictions provide specific biophysically grounded targets for future experimental testing. In total, our results provide novel insight into fundamental biomarkers of innocuous and noxious somatosensory processing.

## 2. Materials and Methods

### 2.1. Median nerve evoked response data interpreted with HNN

Our HNN examination of the MN response focused on interpreting the trial-averaged contralateral MN response in the right hemisphere of a representative human participant (n=1 healthy male human participant, age 39; 111 trials). This dataset was originally published in Sorrentino et al., (2009) and is publicly available with MNE-Python (Gramfort et al., 2014; https://mne.tools/stable/overview/datasets_index.html#somatosensory). The dataset is comprised of the MEG response from 306 sensors to electrical median nerve stimulation of the left wrist at the motor threshold with a uniformly random-sampled interstimulus interval between 7.0 and 9.0 s, as described in the original study, and was collected with written informed consent from the participant and prior approval from the local ethics committee (Sorrentino et al., 2009).

To begin, we performed source analysis on this sensor level data. Given conflicting results in the literature regarding the true number of current dipole sources in and around SI responsible for the early somatosensory evoked field to transcutaneous electrical stimulation of the hand (Ishibashi et al., 2000) and/or median nerve (Allison et al., 1989; Huttunen et al., 2006; Kawamura et al., 1996; Lin et al., 2005, 2003; Peterson et al., 1995; Tiihonen et al., 1989; Wikström et al., 1996), we adopted an approach for source-localizing the MN response that made no a priori assumption about the number of true generators. Using MNE-Python (Gramfort et al., 2013), we fitted a distributed inverse solution by calculating the noise-normalized (i.e., the dynamic statistical parametric mapping [dSPM]) minimum norm estimate inverse solution. We then selected the vertex with maximal cumulative activation over the first 40 ms post-stimulation time window and extracted the time course of activity from the selected vertex from the non-noise-corrected minimum norm estimate (MNE). While this method differs from other methods that fit an equivalent current dipole (ECD) or estimate an average response in a highly active region-of-interest (ROI), our selected vertex represents dipolar current flow constrained to a small, highly responsive region of SI.

### 2.2. Laser evoked response data interpreted with HNN

Our HNN examination of the LE response focused on interpreting the group-averaged (n=107 healthy human participants ages 18-26, 67 of which were female; 80 trials/participant) contralateral LE response computed from ECD modeling that was published in a prior study (Jin et al., 2018). All human participants associated with this dataset provided written informed consent and the experimental procedures were approved by the local ethics committee. As outlined by Jin and colleagues, laser stimulus was applied 40 times to the dorsum of the left and right hands of each participant with a uniformly random-sampled interstimulus interval between 11 and 15 s, and whose laser-beam target was shifted ∼1 cm after each stimulus to avoid overheating of the skin and nociceptor fatigue or sensitization. The energy level of each stimulus was randomly sampled from four values that corresponded to subjective pain ratings of ∼2, ∼4, ∼6, ∼8 on a 0-10 numerical rating scale. The contralateral SI ECD to the left or right hand was fitted to EEG scalp topography as part of a four-dipole model (Jin et al., 2018). Here, we averaged the contralateral SI response across both hemispheres to calculate the cumulative mean EEG evoked response to cutaneous laser stimulation applied to either hand.

### 2.3. HNN computational neural modeling

The HNN software was designed for cellular and circuit level interpretation of source-localized human M/EEG signals and enables one-to-one comparison between simulated and recorded current dipole sources in equal units of nano-ampere-meters (Neymotin et al., 2020). The core of HNN is a pre- constructed and pre-tuned model of a laminated cortical circuit under thalamocortical and cortico-cortical influences that simulates the primary electrical currents (i.e., current source dipoles) generating M/EEG signals from a localized patch of the cortex (Figure 1). HNN’s model was designed to include canonical features of cortical circuitry, including synaptically connected pyramidal neurons and interneurons in supragranular and infragranular layers, and layer specific feedforward and feedback synaptic drive. HNN simulates the primary current sources of M/EEG from the net intracellular current flow across the pyramidal neuron populations in the model, based on the established biophysical origin of these currents (Hämäläinen et al., 1993; Murakami and Okada, 2006; Næss et al., 2021; Okada et al., 1997). In practice, users activate the network by defining patterns of action potentials representing activity in exogenous networks (e.g., thalamus or higher order cortex) that drive the local network through layer specific patterns of excitatory synaptic inputs. Synaptic inputs that drive the network target the dendrites of excitatory pyramidal cells and the somas of inhibitory basket cells. One pathway of input simulates perturbations ascending from the peripheral nervous system, through the lemniscal thalamus to the granular layers (L4), and finally propagating to the proximal dendrites of the pyramidal neurons in supra- and infragranular layers. These inputs are referred to as proximal drive and represent “feedforward” activation of the circuit. The other pathway of input comes from the non-lemniscal thalamus or high-order cortex and synapses in the supragranular layers, targeting the distal dendrites of the pyramidal neurons. These inputs are referred to as distal drive and can represent “feedback” activation of the circuit (Neymotin et al., 2020).

**Figure 1.**
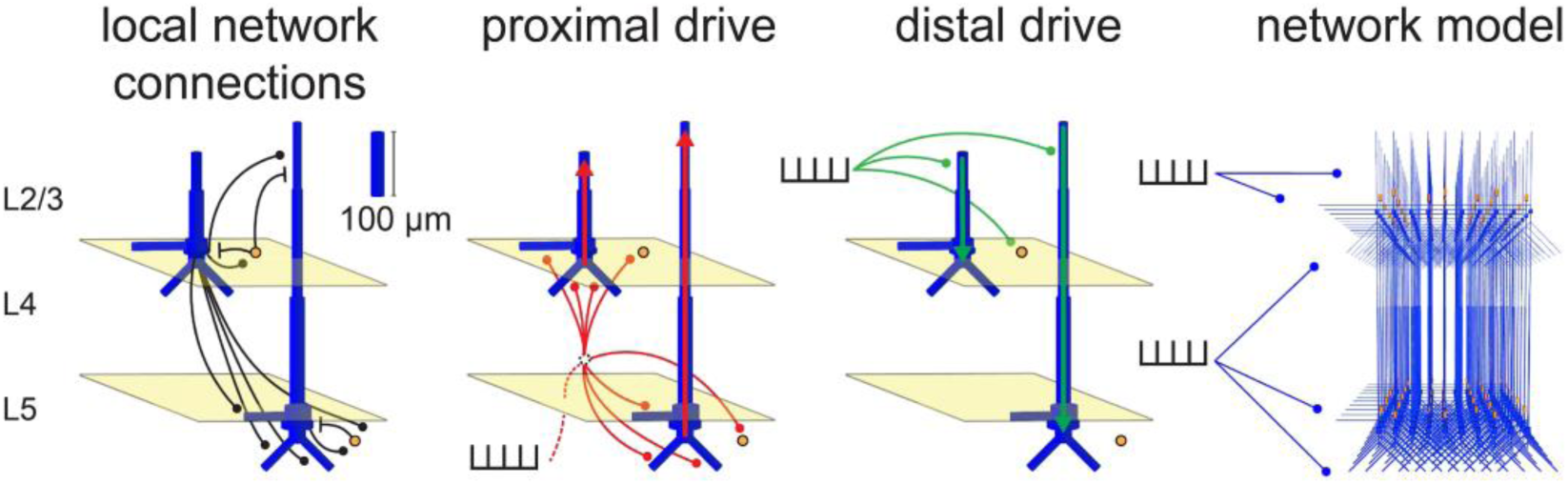
HNN’s foundation is a pre-tuned biophysically principled neocortical network model, under thalamocortical and cortico-cortical drive that simulates primary electrical current dipoles via current flow up and down the long apical dendrites of pyramidal neurons. Local network connections (black) bridge excitatory pyramidal cells (blue) with inhibitory basket cells (yellow circles) and each other. Proximal inputs (red) arrive as feedforward excitatory drive from the lemniscal thalamus and target the granular layer (L4; middle) that propagate directly to the proximal dendrites of the pyramidal neurons in supragranular (L2/3; top) and deep layers (L5; bottom). The initial circuit response of a proximal drive is to push current up the apical dendrites of pyramidal cells. Distal inputs (green) arrive as feedback excitatory drive from the non-lemniscal thalamus or high-order cortex and target the distal portion of the apical dendrite in the supragranular layer. The initial circuit response of a distal drive is to push current down the apical dendrites of pyramidal cells. Inhibitory drive to the circuit and other more complex interactions result from the spatiotemporal dynamics that emerge from basket cell activation. The canonical model network is composed of 35 L2/3 basket cells, 35 L5 basket cells and a 10x10 grid of L2/3 and L5 pyramidal cell pairs that are connected across the grid according to type with exponential synaptic weight fall-off. Adapted from Neymotin et al. (2020).

For both the evoked proximal and distal drive, the synaptic input is generated by simulating a pre- synaptic action potential whose timing on any trial is chosen from a Gaussian distribution with a defined mean time and standard deviation. A *simulation* is composed of multiple trials (here, n=100). Stochasticity across trials occurs through the Gaussian jitter in timing of the proximal and distal drive times. Note that all drives in the present study were synchronously applied to all cells (i.e., one spike time was sampled to supply an excitatory synaptic input to all cells in the network for each drive of a given trial). The HNN model represents synaptic weight as the maximal conductance associated with a given receptor’s IPSC or EPSC. Exogenous proximal and distal drives produce only excitatory synaptic inputs of both fast and slow AMPA/NMDA kinetics onto the pyramidal and inhibitory neurons, as shown in Figure 1. In the local network, pyramidal neurons provide both fast and slow AMPA/NMDA-mediated excitation, and inhibitory neurons provide both fast and slow GABA_A_/GABA_B_-mediated inhibition. Slight adjustments to the local network connection strengths were made from the distributed HNN neocortical template model to prevent runaway firing given the number and types of evoked inputs (Table 1). The same local configuration was used for MN and LE responses.

**Table 1.** Local network parameters of the computational model. Each value represents a synaptic weight (µS) for an excitatory (i.e., AMPA or NMDA) or inhibitory (i.e., GABA_A_ or GABA_B_) connection. Asterisks denote parameters that were initially tuned from the default values distributed with HNN to prevent runaway network excitability and then left constant for all simulations presented in the results of this study.

|  | L2/3 Pyramidal | L5 Pyramidal | L2/3 Basket | L5 Basket |
| --- | --- | --- | --- | --- |
| L2/3 Pyramidal | AMPA: 2e-4*<br>NMDA: 2e-4* | AMPA: 2e-4*<br>NMDA: n/a | AMPA: 5e-4<br>NMDA: n/a | AMPA: 2.5e-4<br>NMDA: n/a |
| L5 Pyramidal | n/a | AMPA: 1e-3*<br>NMDA: 5e-4 | n/a | AMPA: 5e-4<br>NMDA: n/a |
| L2/3 Basket | GABA<sub>A</sub>: 5e-2<br>GABA<sub>B</sub>: 5e-2 | GABA<sub>A</sub>: 2e-3*<br>GABA<sub>B</sub>: n/a | GABA<sub>A</sub>: 2e-2<br>GABA<sub>B</sub>: n/a | n/a |
| L5 Basket | n/a | GABA<sub>A</sub>: 2e-2*<br>GABA<sub>B</sub>: 6e-2* | n/a | GABA<sub>A</sub>: 2e-2<br>GABA<sub>B</sub>: n/a |

The aggregate current dipole is computed by summing the intracellular currents in all pyramidal neurons across both L2/3 and L5 flowing in a direction parallel to the apical dendrites; however, due to the longer length of the L5 pyramidal neurons, the net current dipole response is dominated by the activity in L5. Microcircuit activity, such as cell spiking profiles, can also be visualized leading to detailed predictions on the circuit mechanisms inducing the recorded macroscale current dipole signal. The parameters describing the dynamics of the individual cells, and local and exogenous synaptic interactions have all been set to realistic starting values based on prior studies (Neymotin et al., 2020).

It is important to note that HNN is designed to simulate source-localized current dipoles. The present study uses source-localized data from MEG for the MN response, and EEG for the LE response. While MEG and EEG measure magnetic and electric fields, respectively, they both reflect fields from a primary current dipole source (Hämäläinen et al., 1993), which here is assumed to be originating in SI. In the case of the earliest components of the MN and LE responses, prior studies have shown that MEG and EEG render similar dipole locations orientations in SI, but that EEG produces higher inter-subject variability (Ahmed Mahmutoglu et al., 2022; Hoshiyama and Kakigi, 2001; Zainea et al., 2005). While the absolute magnitude of the MN and LE responses may change from subject to subject, the timing of each major deflection remains relatively stable (Ahmed Mahmutoglu et al., 2022; Ou et al., 2007). Given our emphasis on interpreting stable neural dynamics underlying evoked response deflections, differences between MEG and EEG as well as inter-subject variability should have minimal impact on the results and conclusions discussed herein.

Further details and HNN tutorials can be found at https://hnn.brown.edu/. All code is open-source and provided online at https://github.com/jonescompneurolab/hnn, and further functionality is available at https://github.com/jonescompneurolab/hnn-core. The specific parameter files used here will be distributed upon publication.

### 2.4. Process for hypothesis testing, parameter estimation, and sensitivity analysis in HNN

#### Hypothesis testing in HNN

As described above, HNN is distributed with a pre-constructed and tuned neocortical network model representing a localized patch of cortex that can be perturbed by excitatory synaptic drive through proximal/feedforward and distal/feedback pathways. Due to the large-scale nature of the neocortical model, which has hundreds of parameters, in practice users leave most of the parameters fixed and adjust only a user-defined subset of parameters based on hypotheses motivated by prior studies and literature. The graphical user interface, and quantification of the root mean squared error (RMSE) between simulated and empirical evoked response waveforms, allows users to examine if and how adjustments in these parameters can account for recorded current dipole source data.

Given the HNN neocortical template model (Figure 1, Table 1), all simulations begin by “activating” the network with some type of assumed exogenous drive that depends on the experimental conditions. For sensory evoked responses, as studied here, it is hypothesized based on prior studies that the local network receives a sequence of thalamocortical and/or cortico-cortical drives that occur time-locked to sensory stimulation. In the model, these are simulated via proximal and distal drives. Here, we focussed only on estimating the number, timing and maximal conductance of these drives in order to best account for the recorded data. Initial and several alternative hypotheses on the parameters of these drives are tested to identify configurations that reproduce the best fit to data (i.e., smallest RMSE). The best fit model provides targeted predictions on the multiscale mechanisms creating the current dipole source signal that can guide further follow up testing with invasive recordings or other imaging modalities (e.g., see (Bonaiuto et al., 2021; Sherman et al., 2016)).

Since the HNN neocortical template model contains only a 10x10 grid of pyramidal cells, which is in most cases significantly smaller than the actual number of cells contributing to a recorded current dipole, the second step in the HNN simulation is to adjust the dipole “scaling factor” that multiplies the amplitude of the simulated net current dipole by a value that makes the amplitude of the simulated and recorded signals comparable (see further details on the HNN website and Neymotin et al. (2020). Here, the dipole scaling factor was set to 10, 40, and 2500 for the TE, MN, and LE responses, respectively. Further, all simulated dipole trials were smoothed by convolution with a Hamming window (5 ms width during parameter tuning/optimization to prevent overfitting, 20 ms for the final product) in order to better match the signal-to-noise ratio of empirical current dipoles recorded from real neural networks that are larger and more diverse.

#### Parameter estimation

The process for parameter estimation begins by starting with a hypothesized sequence of drives, using the drive parameters distributed with the software, and manually “hand-tuning” the parameters defining the timing and strength of these drives to get an initial close representation to the current dipole waveform, i.e., small RMSE between the empirical current dipole and simulated trial-mean current dipole. These simulations are referred to as “pre-optimization” and are characterized by a pre-optimization parameter set (pre-OPS). Once an initial close representation to the data is found, automated parameter estimation algorithms distributed with the software can be used to estimate parameters (within a defined range) that produce the smallest RMSE between the simulated and recorded current dipoles (i.e., the “best fit” to the data). These simulations are referred to as “post-optimization” and are characterized by a post-optimization parameter set (post-OPS). Here, optimization routines were configured to explore all exogenous proximal and distal drive-related parameters and run 3 iterations per parameter in each optimization stage (see Figures 6, 8, and 10 for a complete list of parameters). Each iteration consisted of a full simulation (i.e., with n=100 trials, 170 and 300 ms/trial for the MN and LE responses, respectively) except for any simulation with 5 or more drives, which given its computational cost, consisted of 50 trials/simulation to reduce run-time.

Of note, the pre-tuned HNN model and above described process for hypothesis testing and parameter estimation has been successfully applied in several prior studies of sensory evoked responses (Jones et al., 2009, 2007; Kohl et al., 2021; Law et al., 2021; Sliva et al., 2018).

#### Sensitivity analysis

Once an optimized solution was found, parameter sensitivity analysis was performed on an identified subset of key parameters to examine how each contributed to variance in the simulated current dipole. The chosen parameters-of-interest for the optimized MN response (Figure 4d) were informed by a prior sensitivity analysis on the HNN model that showed that several of the evoked proximal and distal drive parameters accounted for the largest amount of variance in the TE current dipole signal, as defined by their Sobol sensitivity indices (Neymotin et al., 2020). Parameters of the evoked proximal drives included their mean drive time and the maximal postsynaptic conductance of the NMDA currents only, specifically L2_pyramidal_nmda, L5_basket_nmda, L5_pyramidal_nmda. Parameters of the distal drive included their mean drive time and the maximal postsynaptic conductions of both AMPA and NMDA currents, specifically L2_basket_ampa, L2_pyramidal_nmda, and L5_pyramidal_nmda. For the optimized LE response (Figure 9d), parameters were selected using knowledge from simulation exploration such that the selected subset produced the greatest variance in the signal. Parameters of the first evoked proximal drive included the mean drive time and the maximal postsynaptic conductance of AMPA and NMDA, specifically L2_basket_ampa, L2_pyramidal_ampa, L5_basket_nmda. Parameters of the second evoked proximal drive included the mean drive time and AMPA and NMDA currents onto the pyramidal neurons, specifically L2_pyramidal_ampa, L5_pyramidal_ampa, L5_pyramidal_nmda. Parameters of the distal drive included the mean drive times of the burst of inputs and their inter-drive-interval (IDI), as well as the maximal postsynaptic conductance of AMPA and NMDA currents, specifically L2_basket_ampa, L2_pyramidal_ampa, L2_basket_nmda. The sensitivity analysis was conducted using batch scripts and the Python application programming interface for HNN, HNN-core (https://github.com/jonescompneurolab/hnn-core), by sweeping through a range of each identified parameter, while keeping all other parameters fixed, and calculating the net standard deviation contributed by that parameter and the cumulative increase across all parameters, at each point in time (Figure 4d and Figure 9d. Each parameter sweep consisted of 50 simulations where the parameter-of-interest was sampled from a uniform distribution while the other parameters remained constant. Boundary conditions were applied to each varied parameter as follows: synaptic weight parameters were sampled from 10^-5^-10^-2^ µS on a log_10_ scale while drive timing parameters were sampled over (post-OPS)±4 ms for the MN case and (post-OPS)±10 ms for the LE case.

### 2.5. Time-frequency response analysis

The average time-frequency response (TFR) was calculated from 3-100 Hz across simulated aggregate current dipole trials by computing each trial’s TFR through convolution of the zero-padded time course with an *m*=*f*/3 cycle Morlet wavelet of the form \[w(t,f)=A\cdot exp\left(\frac{-t^2}{2\sigma^2}\right)\cdot exp(i2\pi ft)\], where \[A=\left(\frac{1}{2}\sum_{t}\vert w(t,f)\vert ^2\right)^{-1/2}\] and \[\sigma=\frac{m}{2\pi f}\] for a given frequency value *f*. To preserve high-frequency components and minimize edge effects in the TFR calculation of each time course, trials were smoothed with a 5 ms-wide Hamming window and run for a longer length of time for a total of 600 ms. TFR trials were then cropped at t=300 ms. Given that the TFR was only calculated for the LE response simulations which maintained a current dipole close to zero for the first ∼100 ms, edge effects at the beginning of each time course were negligible.

## 3. Results

### 3.1. MN and LE responses have robust temporal profiles comprised of clear waveform features that were the targets of mechanistic investigation with HNN

The MN evoked response has been established as a robust and reliable signal that can be source-localized to a current dipole in SI, albeit differences in waveform features emerge due to different recording methods and individualized responses. Despite this variability, there are clear consistencies in the timing and orientation of the response waveform, which historically served as the foundation for the peak-latency labeling nomenclature (e.g., N20, P30, etc). Before studying the cell and circuit mechanisms generating MN responses with HNN, we first defined a set of waveform commonalities across representative amplitude-normalized MN responses reported in prior studies (Figure 2a). Each of the example MN response traces in Figure 2a showed maximal activity from 20-150 ms post-stimulus. There are four common features referred to as N1, P1, P2/prolonged depression, and rebound that we aimed to study with HNN. Specifically,

**Figure 2.**
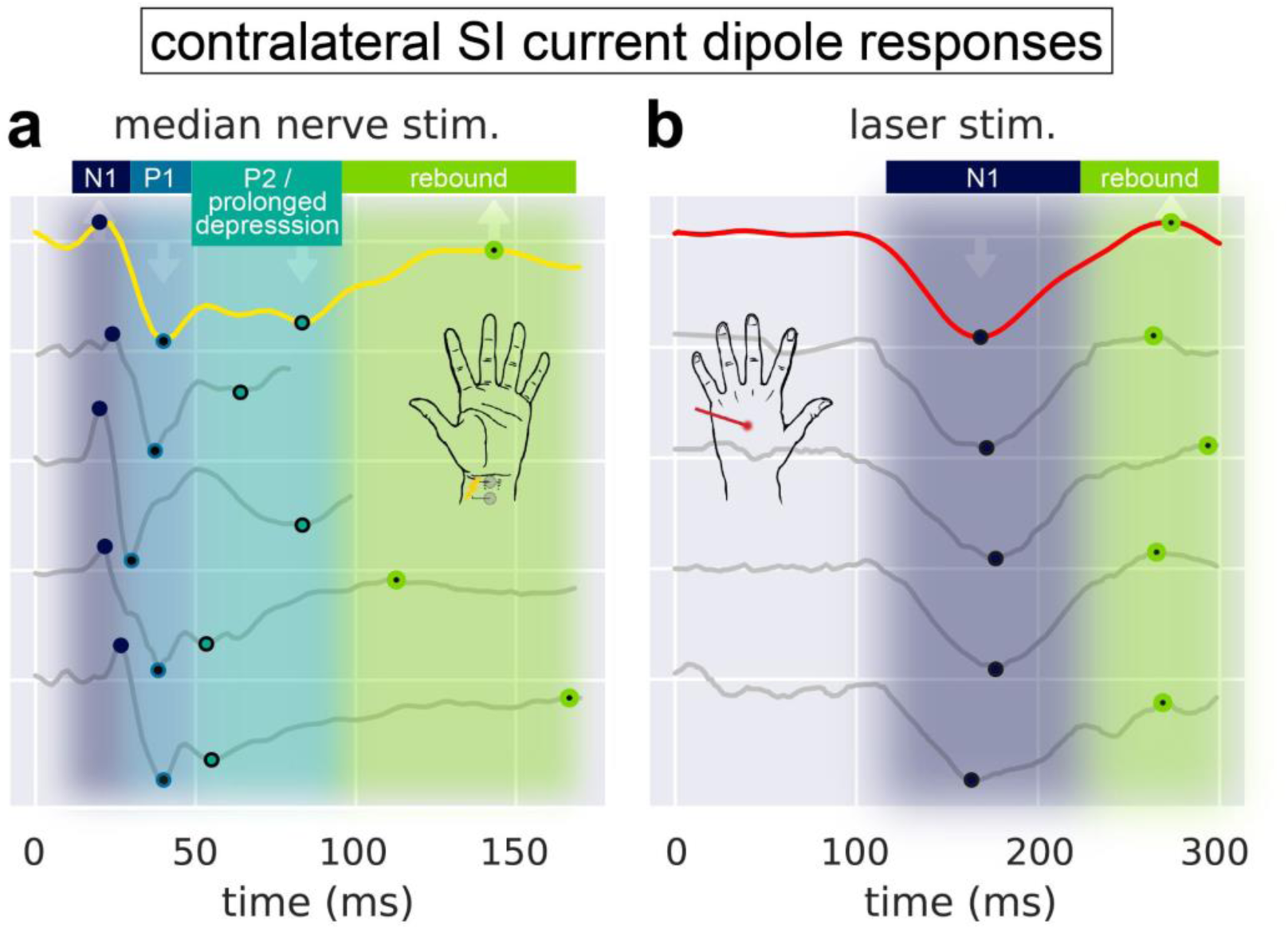
Primary deflections of the MN and LE responses are robust, reproducible, and conserved across multiple M/EEG experiments. (a) The exemplar MN current dipole response analyzed in this study (yellow; MEG inverse solution of n=1 human subject) exhibits three deflections (N1, P1, and P2/prolonged depression) followed by a rebound that are broadly conserved across other studies (grey). Example MN current dipole responses have been reproduced here from prior studies and oriented so that their first major deflection points upward, showing consistent peak alignment despite different underlying experimental conditions: EEG with n=1 (Zainea et al., 2005); MEG with n=13 (Inui et al., 2004); MEG with n=1 (Forss et al., 2001), and MEG with n=1 (Lin et al., 2003). (b) Same as for the left side, except that the LE current dipole response (red; EEG inverse solution of n=107 human subjects) contains only one deflection (N1) followed by a small rebound peak. Example LE current dipole responses have been reproduced here from prior studies and oriented so that their first major deflection points downward, showing consistent peak alignment despite different underlying experimental conditions: EEG with n=12 (Ahmed Mahmutoglu et al., 2022), MEG with n=12 (Ahmed Mahmutoglu et al., 2022), EEG with n=10 (Schlereth et al., 2003), and MEG with n=1 (Raij et al., 2003). Each waveform in (a) and (b) has been normalized by the magnitude of its most prominent deflection. Numerical data for the example current dipole responses (grey traces) were extracted using the WebPlotDigitizer application (https://automeris.io/WebPlotDigitizer/).

1. The N1 occurs near 20 ms and its magnitude is small compared to the neighboring P1 deflection.
2. The P1 occurs between 35-40 ms and is the sharpest and most prominent deflection.
3. After the P1, there is a prolonged depression of duration between 50-100 ms (P2/Prolonged depression).
4. After the prolonged depression, there is a rebound to baseline.

The source-localized LE response is also highly consistent across prior studies, with distinguishing features compared to the MN response (Figure 2b). The LE response emerges more slowly such that the majority of its activity occurs from 100-300 ms post-stimulus. There are two clear commonalities that we aimed to study with HNN. Specifically,

1. The N1 is a large and prolonged deflection that emerges between 100-200 ms.
2. After the N1, there is a smaller magnitude rebound up to 300 ms.

The goal of HNN modeling was to study the circuit mechanisms of the waveform commonalities despite variability across studies. The exemplar yellow MN and the red LE responses in Figure 2 (each representing a trial-averaged, source-localized evoked response) were the waveforms used for circuit level investigation with HNN below, and a subsequent parameter sensitivity analysis showed how variability around the common features may emerge. It is important to note that HNN is designed to simulate source-localized current dipoles with directly comparable units of nAm. Further, the present study uses source-localized data from MEG for the MN response, and EEG for the LE response. While MEG and EEG measure magnetic and electric fields, respectively, they both reflect fields from primary current sources (Hämäläinen et al., 1993), which in this study have been isolated to originate in SI. In the case of the earliest components of the MN and LE responses, several studies have shown that MEG and EEG render similar dipole locations orientations in SI, but that EEG produces higher inter-subject variability (Ahmed Mahmutoglu et al., 2022; Hoshiyama and Kakigi, 2001; Zainea et al., 2005). While the absolute magnitude of the MN and LE responses may change from subject to subject, the timing of each major deflection remains relatively stable (Ahmed Mahmutoglu et al., 2022; Ou et al., 2007). Given our emphasis on interpreting robust commonalities in the evoked response deflections of the MN and LE cases, respectively, differences between MEG and EEG as well as inter-subject variability should have minimal impact on the results and conclusions discussed herein.

### 3.2. The earliest MN and LE N1 deflections represent current flow in opposite directions across the neocortical layers

A first step in interpreting the circuit mechanisms generating the source-localized MN and LE responses with HNN is to determine the direction of electrical current flow in relation to the neocortical layers across time. This requires the application of inverse solution methods explicitly designed to estimate the location and directionality of the primary currents at specific points of time. Each of the example responses in Figure 2 represents a source-localized current dipole in SI in which information about the direction of current flow into or out of the cortex can be inferred. Notably, traces in Figure 2 have been oriented such that that positive deflections correspond to dipole currents directed from deep to superficial layers (i.e., out of the cortex) while negative deflections correspond to currents directed from superficial to deep layers (i.e., into the cortex).

More specifically, the ∼20 ms MN N1 corresponds to current flow out of the cortex, while the ∼40 ms MN P1 corresponds to current into the cortex. This is explicitly shown in Figure 3 where we have applied a minimum norm estimation method to the sensor-level MN response data in order to extract the source-localized and oriented response in Figure 3b (also shown in Figure 2, yellow trace). Figure 3a shows the location and absolute magnitude of the initial response from MN stimulation averaged over the first 40 ms, and Figure 3b shows the dipole moment time course at the vertex of peak activation. The orientation and forward (i.e., posterior-to-anterior) direction at the time of the MN N1 dipole moment at this vertex reflects current flow traveling from deep to superficial layers (out of the cortex, Figure 3c). Conversely, the MN P1 at 40 ms, reflects backward (i.e., anterior-to-posterior) current flow traveling from superficial to deep layers (into the cortex, Figure 3d). The cross-sectional cut-out of the central sulcus (CS) and postcentral gyrus rendered in Figure 3e illustrates how the inferred laminar orientation of a dipole moment deflection depends on the location within the folds of SI.

**Figure 3.**
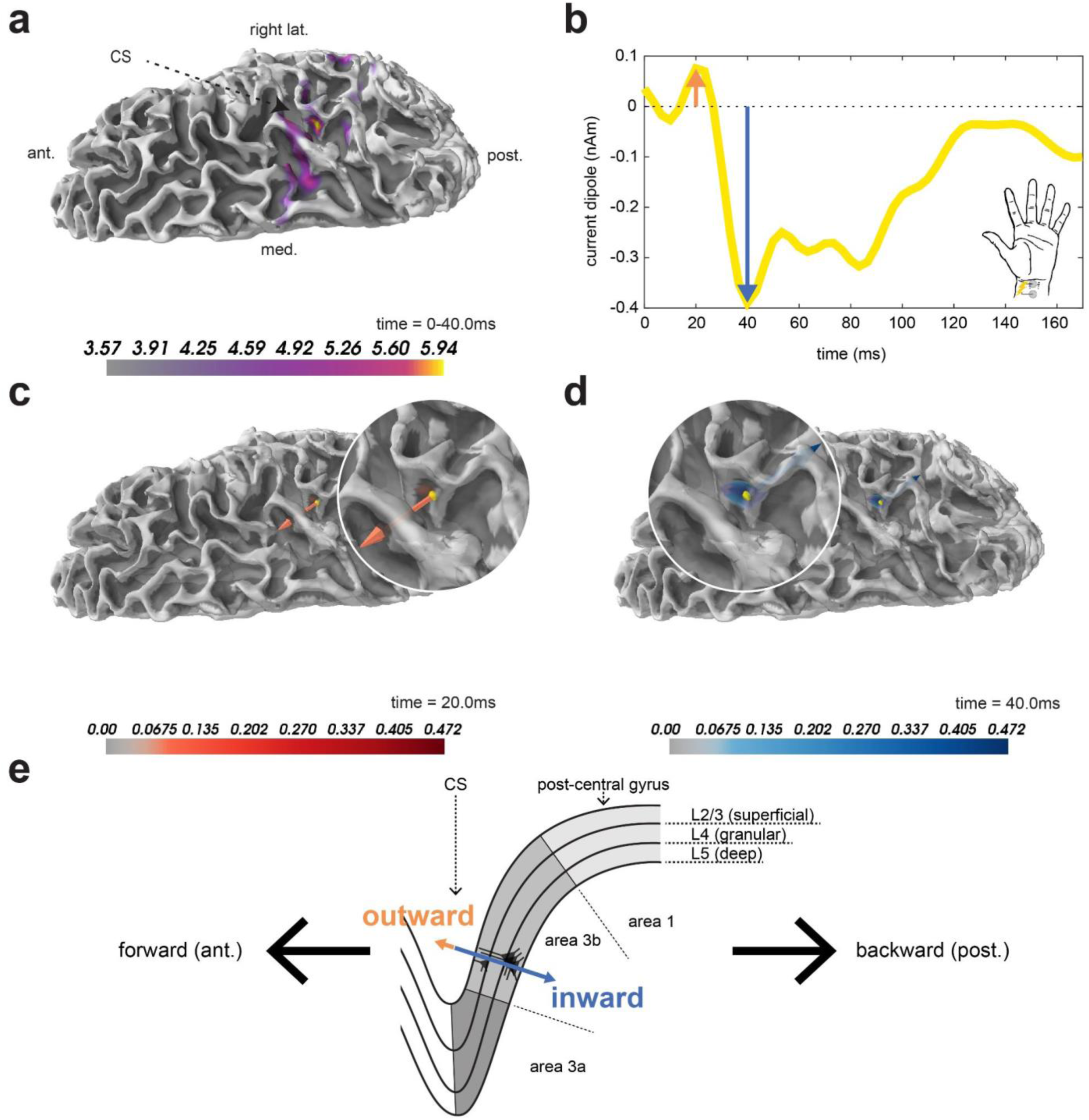
Waveform deflections in M/EEG source-space correspond to sequential, orientation-specific activation of a contralateral median nerve-responsive cortical dipole. (a) Dorsal view of the right hemisphere’s average dynamic statistical parametric map (dSPM) of the minimum norm estimate (MNE) over the first 40 ms post-stimulation. To isolate a vertex-of-interest (VOI), the single most active early-latency vertex was selected from the dSPM and shown in yellow. Note that maximal dSPM activity occurs in the hand representation of area 3b on the posterior wall of the central sulcus (CS). (b) The MN response found by selecting the MNE time course at the VOI. (c) Direction and orientation of the current dipole at 20.0 ms. (d) Same as in (c), except at 40.0 ms. (e) Cartoon schematic demonstrating how the location and mesoscopic direction of the current dipole correspond to laminar orientation and direction at each deflection of the dipole time course. The first major deflection of the MN response at 20.0 ms represents forward current flow oriented *out* of the cortex (i.e., from deep to superficial layers; outward currents). The second major deflection at 40.0 ms represents backward current flow oriented *into* the cortex (i.e., from superficial to deep layers; inward currents).

The outward directionality of the MN N1 has been confirmed in several studies, as detailed in Table 2, and is consistent with prior invasive recordings of MN responses with cortical surface electrocorticography (ECoG) and cortical depth electrodes in human neurosurgery and epilepsy patients (Allison et al., 1989). The nomenclature of “negative” for the “N1” may seem counterintuitive at the source level since the current dipole source is actually directed in the “positive” direction (i.e., out of the cortex), but the negative nomenclature comes from the fact that it corresponds to a current sink in the posterior (referenced to anterior) cortex as measured from EEG sensors. More specifically, the earliest electrical field of somatosensory evoked potentials at the posterior sensor minus anterior sensor is negative (Deiber et al. 1986).

**Tabel 2.** Directionality of the contralateral SI dipole current from MN stimulation at the time of the N1 deflection in prior publications. Stimulus was applied to the median nerve of the hand and its evoked response measured in the relative contralateral hemisphere of the cortex. *Example evoked response data from this study is depicted in Figure 2a. ^†^This study delivered transcutaneous electrical stimulation to the dorsum of the right hand rather than median nerve stimulation. ^††^This study delivered tactile stimulation to the first digit of the hand rather than median nerve stimulation. Left hemisphere, LH; right hemisphere, RH; standard error of the mean, SEM; standard deviation, SD.

| Direction and inferred laminar orientation of the contralateral MN response dipole at its N1 deflection in prior publications |  |  |  |  |  |  |
| --- | --- | --- | --- | --- | --- | --- |
| Publication | Measurement modality | Group-level/single subj. dipole fit | Peak time (mean ±SEM) | Dipole location (Brodmann area) | Primary dipole direction | Inferred laminar dipole orientation |
| Inui et al., 2004*† | MEG | Single (n=13) | RH: 20.8 | 3b | forward | outward |
| Jones et al., 2007*†† | MEG | Single (n=7) | LH: 23.3 | 3b | forward | outward |
| Kanno et al., 2003 | MEG | Single (n=14) | 19.8±0.2 | 3b | forward | outward |
| Lin et al., 2003* | MEG | Single (n=13) | RH: 21.7±0.4 | 3b | forward | outward |
| Lin et al., 2005 | MEG | Single (n=17) | RH: 22.1±0.2 | 3b | forward | outward |
| Tiihonen et al., 1989 | MEG | Single (n=6) | RH: 19.7±1.8 (mean±SD) | 3b | forward | outward |
| Wikstrom et al., 1996 | MEG | Single (n=9) | RH: 21.0±1.6 (mean±SD) | 3b | forward | outward |

The LE response is less well studied at the source level. Unlike the MN N1 response, which is consistently localized to area 3b, the precise localization of the initial LE N1 response is less consistent, with some studies localizing closer to the anterior wall of the postcentral gyrus (e.g., areas 3a/3b), other studies localizing near the crown and posterior wall of the gyrus (e.g., areas 1 and 2), and some studies lacking fine spatial resolution altogether. Invasive recordings in non-human primates support the crown of the postcentral gyrus (nearer to area 1) as being a primary target of nociceptive projections to SI (Kenshalo et al., 2000; Vierck et al., 2013), however the orientation of the dipole current across the cortical layers has not previously been clearly delineated. The direction of the current dipole at the time of the primary LE N1 deflection can be visualized overlayed on structural MRI images and is summarized in Table 3 for each study. Further, based on the location of the maximal activation along the postcentral gyrus reported in several of these studies, we can infer the current direction into or out of the cortex using the same coordinate frame of reference as depicted in Figure 3e. The majority of the studies suggest that the LE N1 current dipole is directed into the cortex, suggesting current flow down the pyramidal neuron dendrites. Importantly, this is due to a different average source location on the postcentral gyrus and has an opposite current dipole direction compared to the MN N1 response, despite having a similar negative voltage potential at parietal EEG electrodes (hence the negative nomenclature). Here, we have aligned the waveforms so that peaks representing inward currents have negative dipole current values (nAm). As such, the MN N1 has a positive value, while the LE N1 has a negative value.

**Tabel 3.** Directionality of the contralateral SI dipole current from laser stimulation at the time of the N1 deflection in prior publications. Stimulus was applied to the dorsum of the hand and its evoked response measured in the relative contralateral hemisphere of the cortex. *Example evoked response data from this study is depicted in Figure 2b. ^†^This study delivered noxious intraepidermal electrical stimulation to the dorsum of the hand rather than cutaneous laser stimulation. LH, left hemisphere; RH, right hemisphere; standard error of the mean, SEM.

| Direction and inferred laminar orientation of the contralateral LE response dipole at its N1 deflection in prior publications |  |  |  |  |  |  |
| --- | --- | --- | --- | --- | --- | --- |
| Publication | Measurement modality | Group-level/single subj. dipole fit | Peak time (mean ±SEM) | Dipole location (Brodmann area) | Primary dipole direction | Inferred laminar dipole orientation |
| Baumgärtner et al., 2011 | intracranial EEG grid | Single (n=2) | LH: 138<br>RH: 141 | 3a/3b | backward | inward |
| Jin et al., 2018* | EEG | Group (n=107) | LH: 165<br>RH: 170 | non-specific SI | downward/<br>forward | inward |
| Omori et al., 2013† | MEG | Single (n=4) | RH: 169±12 | 3b/1 | backward | inward |
| Ploner et al., 1999 | MEG | Single (n=6) | 174±3 | 3b/1 | backward | inward |
| Raij et al., 2003* | MEG | Single (n=8) | RH: 160-195 | non-specific SI | backward | inconclusive |
| Schlereth et al., 2003* | EEG | Single (n=10) | 180±7 | 1/2 | backward | outward |
| Valentini et al., 2012 | EEG | Group and single (n=35) | LH: 156<br>RH: 157 | non-specific SI | downward/<br>forward | inward |

The differences in the deflections of the respective MN and LE responses indicate distinct underlying mechanisms of generation. We next applied the HNN neural modeling software tool to predict the detailed cell and network level activity generating the robust features of each signal.

### 3.3. HNN predicts the MN response is generated from a specific sequence of exogenous drive similar to that of the TE response, with an additional distal/feedback drive at ∼30 ms

As detailed further in the Methods, HNN is designed to be a hypothesis development and testing tool. The core of HNN is a pre-constructed, pre-tuned model of a canonical neocortical circuit with layer specific activation patterns representing thalamocortical and cortico-cortical excitatory synaptic drive (i.e., proximal and distal drive as shown in Figure 1). HNN simulates a macroscale current dipole in units directly comparable to source-localized data (in units of nAm) while also providing information about the microcircuit activity such as the contribution of different cortical layers, and cell specific spiking responses. In a simulation experiment, most of the parameters in the network are fixed while a targeted set of parameters is estimated (manually and/or algorithmically) based on user defined hypotheses until a close fit to the recorded current dipole data is obtained. In the case of sensory evoked responses, as studied here, it is hypothesized that the local network receives a sequence of thalamocortical and/or cortico-cortical drives that occurs time-locked to sensory stimulation. We focussed only on estimating the parameters of drives to values that could best account for the recorded source-localized data.

We began our investigation with the observation that the SI MN response has a similar waveform shape to the SI TE response to brief stimulation of the fingertip (compare solid yellow and dashed yellow traces, Figure 4a), whose mechanisms of generation have been extensively studied with HNN (Jones et al., 2009, 2007; Law et al., 2021; Neymotin et al., 2020; Sliva et al., 2018). The four MN features we aimed to capture with HNN, as described above, all occur for the TE response (i.e., the N1, P1, P2/prolonged depression, and rebound, as indicated with shading in Figure 4a), albeit with variable amplitude and duration. The most notable differences are that the MN P1 (∼30 ms) has significantly larger magnitude, the P2/prolonged depression is of longer duration, and the rebound to baseline is less pronounced. Given the similarity between the MN and TE responses, our initial hypothesis was that the sequence of exogenous drive previously shown to reproduce the TE response (Jones et al., 2009, 2007; Law et al., 2021; Neymotin et al., 2020; Sliva et al., 2018), could also generate the main features of the MN response. This sequence is depicted in the histograms at the top of Figure 4a, and consisted of a proximal drive at t=19.6±2.5 ms (mean±SD), distal drive at t=65.7±3.8 ms, and a re-emergent proximal drive at t=90.5±10.4 ms. It was based on empirical studies that suggested the tactile stimulation is first relayed in a feedforward manner from the thalamus to SI at ∼20 ms, followed by feedback input to SI at ∼65 ms, followed by re-emerge feedforward input at ∼90 ms that emerges as part of a thalamocortical loop of activation. The parameters of defining each drive were set to previously published values, as distributed with HNN. The resulting simulated current dipole response (mean n=100 trials, black trace, Figure 4a) produced a close representation to the empirical TE response (dotted yellow trace). Variability across individual trials (grey traces) occurs from the stochastic nature of the timing of the drives, which are chosen from a Gaussian distribution (see Methods).

**Figure 4.**
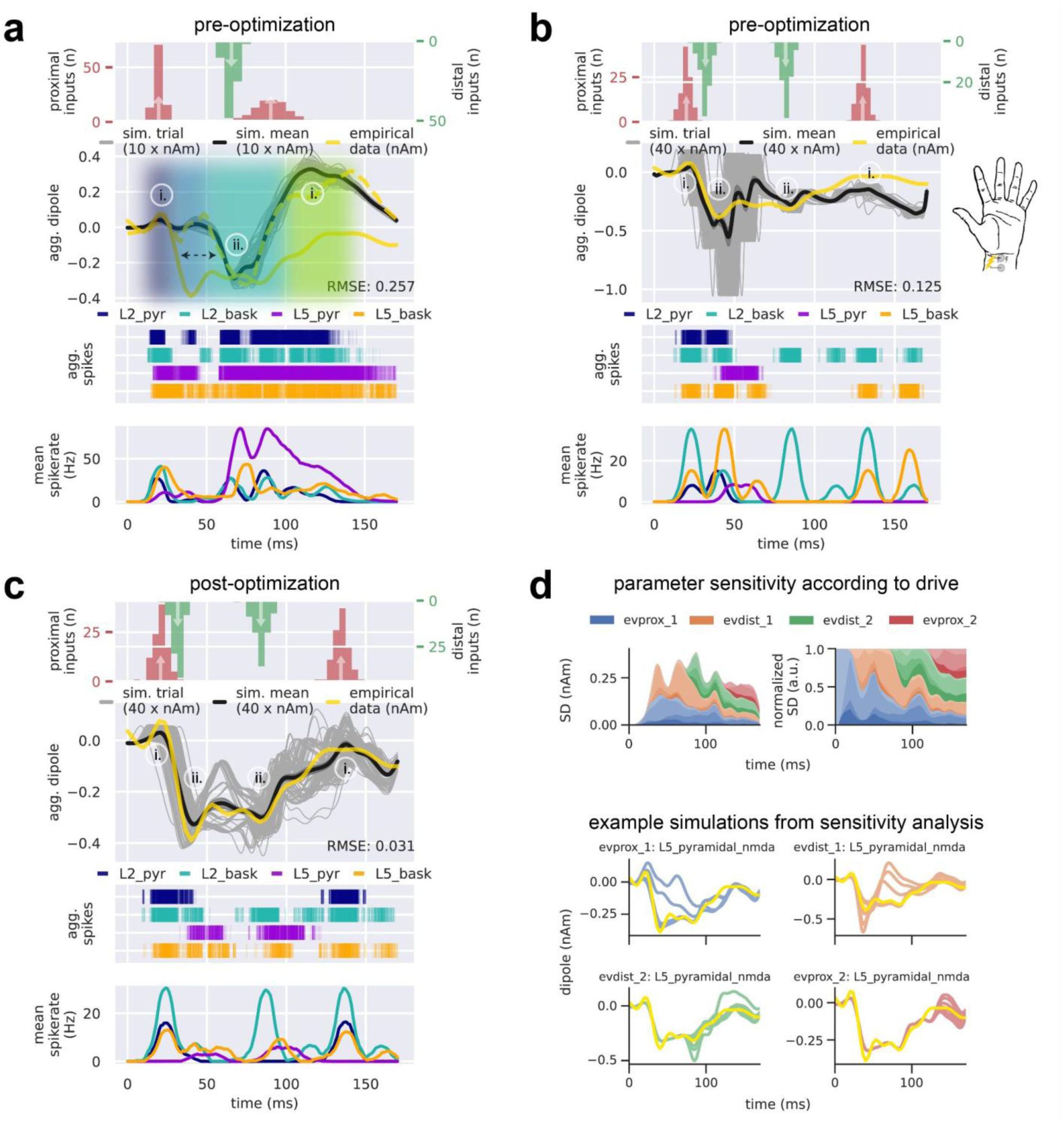
The sequence of outward and inward current flow in the MN response requires a sequence of four drives that begins with an initial proximal drive immediately followed by a strong distal drive. From top to bottom, panels (a-c) each contain the input drive histogram across all 100 trials (proximal, red; distal, green), aggregate simulated dipole (individual trials, grey traces n=100; trial-mean, black trace) alongside the empirical MN response (yellow solid trace), aggregate spike raster for all neurons over all trials according to type (L2 pyramidal, navy; L2 basket, cyan; L5 pyramidal, violet; L5 basket, orange), and population-mean spikerate (i.e., the average spikerate of a neuron of a given type). (a) The simulated suprathreshold TE response explored in prior work (Neymotin et al., 2020) matches the empirical TE response (yellow dashed trace) by producing a prominent negative deflection at ∼65 ms; however, it fails to produce the earliest MN P1 deflection. Note that the TE response simulation uses non-synchronous driving inputs (i.e., drive spikes that are distributed across both time and cells for a given trial). (b) With the addition of a manually-tuned distal drive at t=32.0±3.0 ms, the early upward-downward dynamic characteristic of the MN N1 and P1 is simulated. (c) After parameter optimization of the drive sequence in (b), all deflections and temporal features of the empirical MN response are present and improved upon in the simulated response. (d) Sensitivity analysis of selected parameters belonging to each of the four evoked drives (evprox_1, evprox_2,evdist_1, and evdist_2) illustrates the prominent and differential role respective drives have in controlling the variance of the simulated current dipole waveform at each point in time. Top left: net standard deviation (SD) of the simulated current dipole contributed by each parameter at each time point. Top right: normalized net SD at each timepoint. Parameters of the proximal drives included their mean drive time and the maximal postsynaptic conductance of the NMDA currents only, namely L2_pyramidal_nmda, L5_basket_nmda, L5_pyramidal_nmda. Parameters of the distal drive included their mean drive time and the maximal postsynaptic conductions of both the NMDA and AMPA currents, namely L2_basket_ampa, L2_pyramidal_nmda, and L5_pyramidal_nmda. Bottom: example simulations from the sensitivity analysis showing how changes in the maximal NMDA conductance of each drive impacts the MN response shape, while keeping all other parameters fixed.

Next, we summarize how this drive sequence generated outward and inward current flow that corresponds to the positive and negative deflections in the recorded TE signal, and then describe how we expanded from our initial hypotheses to better account for the notable differences in MN waveform. The first proximal drive at ∼20 ms created excitatory synaptic events that led to depolarization and somatic spiking in the L2/3 and L5 pyramidal neurons. Spike histograms for each cell class across all 100 trials is shown under the current dipole trace in Figure 4a. The somatic action potentials generated strong and fast back-propagation of current up the apical dendrites creating a net outward current flow that accounts for the first positive N1 peak. This peak was dominated by activity in L2/3 (see layer specific responses in Supplementary Figure 1). The subsequent small negative P1 deflection emerged from slightly delayed activation of the interneurons that inhibit the pyramidal neuron somas and pull current flow down the apical dendrites. The second slightly positive peak at ∼50 ms (not labeled) reflected continued spiking of the L5 pyramidal neuron near 40 ms. The subsequent distal drive at ∼65 ms, excited the distal apical dendrite of the L2/3 and L5 pyramidal neurons and pushed inward current flow down the apical dendrite to create the P2 deflection. Additionally, inhibitory cell spiking induced hyperpolarizing currents at the pyramidal somas, pulling current downward (i.e., inward) toward the soma. Subsequent spiking in the L2/3 and L5 pyramidal neurons, together with the re-emergent proximal input at ∼90 ms, induced current flow back up the apical dendrites to create the final outward positive deflection near 100 ms.

In summary, our initial hypothesis was that the sequence of drives that created the TE could reproduce the main features of the MN response (Figure 5a). These simulations reproduced prior published results and recapitulated that outward currents (positive deflections) were created by proximal drive that pushed depolarizing excitatory currents up the pyramidal neuron apical dendrites and/or the backpropagation of somatic action potentials up the dendrites. Inhibition on the distal dendrites of L5 pyramidal neurons also contributed to outward currents during the period of the rebound. Inward currents (negative deflections) were created by distal drive that pushed depolarizing excitatory currents down the pyramidal neuron apical dendrites, facilitated by dendritic calcium activity, and/or somatic inhibition that pulls hyperpolarizing current flow down the apical dendrites.

**Figure 5.**
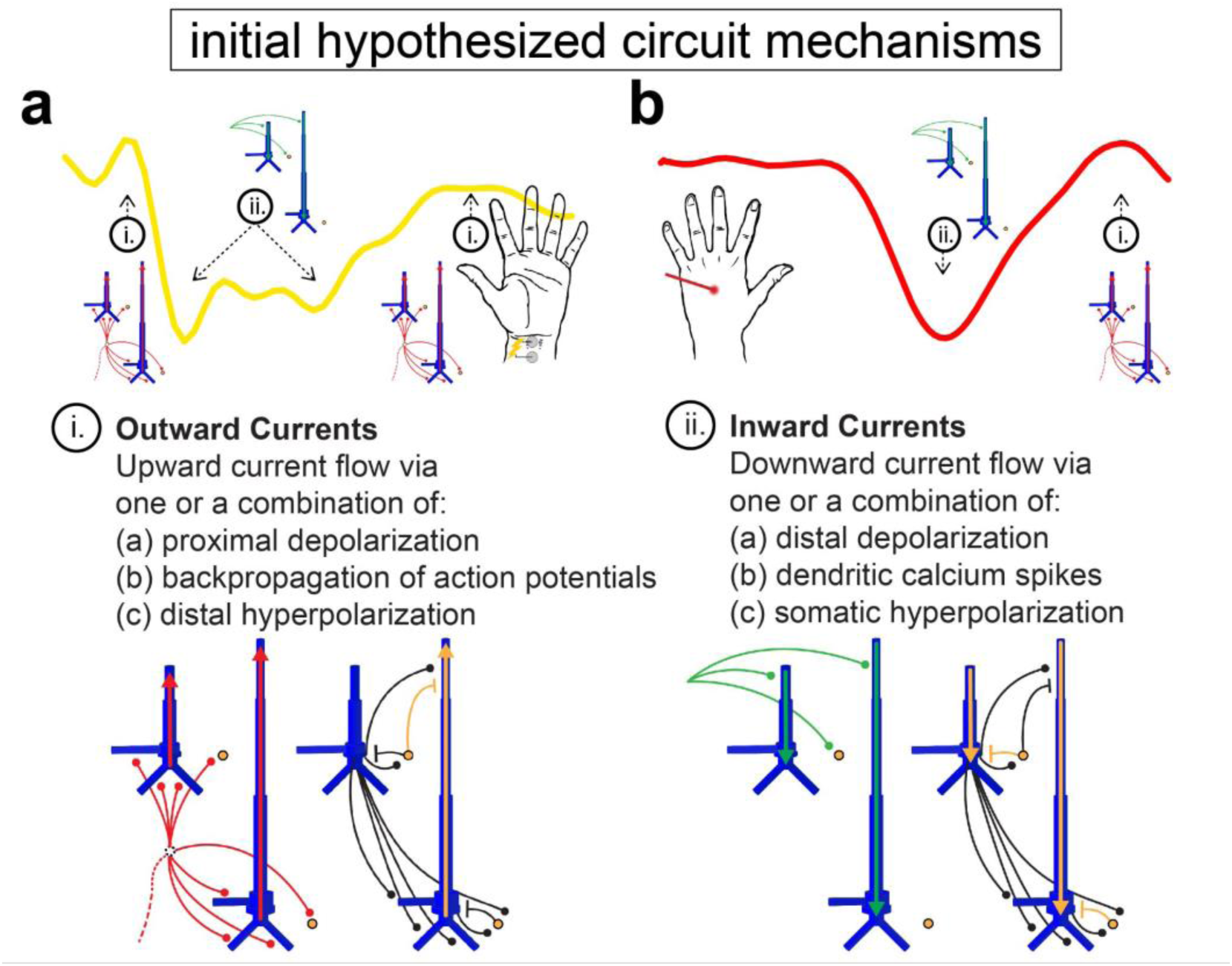
Initial hypothesized sequence of drives and induced circuit mechanisms creating the MN and LE responses. Knowledge obtained from prior studies on the exogenous drives and local network dynamics that can generate outward and inward currents (i.e., positive and negative deflections) led to initial hypothesized circuit mechanisms generating the MN (a) and LE (b) responses.

While the sequence of drives reproduced several of the main features of the MN response, it did not capture the significantly larger magnitude of the MN P1 (∼30 ms), the longer duration of P2/prolonged depression, or the less pronounced return to baseline. We thus expanded our initial hypotheses to try to account for these differences. We first noted that the downward slope from the MN N1 to P1 deflection was consistent with the downward slope to P2 deflection in the simulated TE response (Figure 4a black dashed arrow). Given that the distal drive at ∼65 ms was sufficient to create this steep slope and large negative deflection in the TE response, we considered the effect of adding an additional earlier distal drive at ∼30 ms, hypothesizing that it would similarly create the prominent MN P1. The parameters of this new drive were at first hand-tuned starting with the values from the TE response (Figure 4b; see Figure 6a,b for parameters corresponding to the TE and MN response simulations, respectively). Indeed, combined with the ∼20 ms proximal drive, adding a distal drive at ∼30 ms, recreated a close representation to the MN N1 and P1 deflections. Somatic inhibition also contributed to the prominent P1 generation, where prolonged activation of inhibitory cells in both L2/3 and L5 from 40-50 ms continued to pull current flow down the dendrites and the average current dipole remained negative (see also layer specific responses Supplementary Figure 1b). Some L5 pyramidal neurons were excited enough to spike, and their action potentials induced current flow up the dendrites, creating a slight positive rebound at ∼60-70 ms but never reaching baseline.

**Figure 6.**
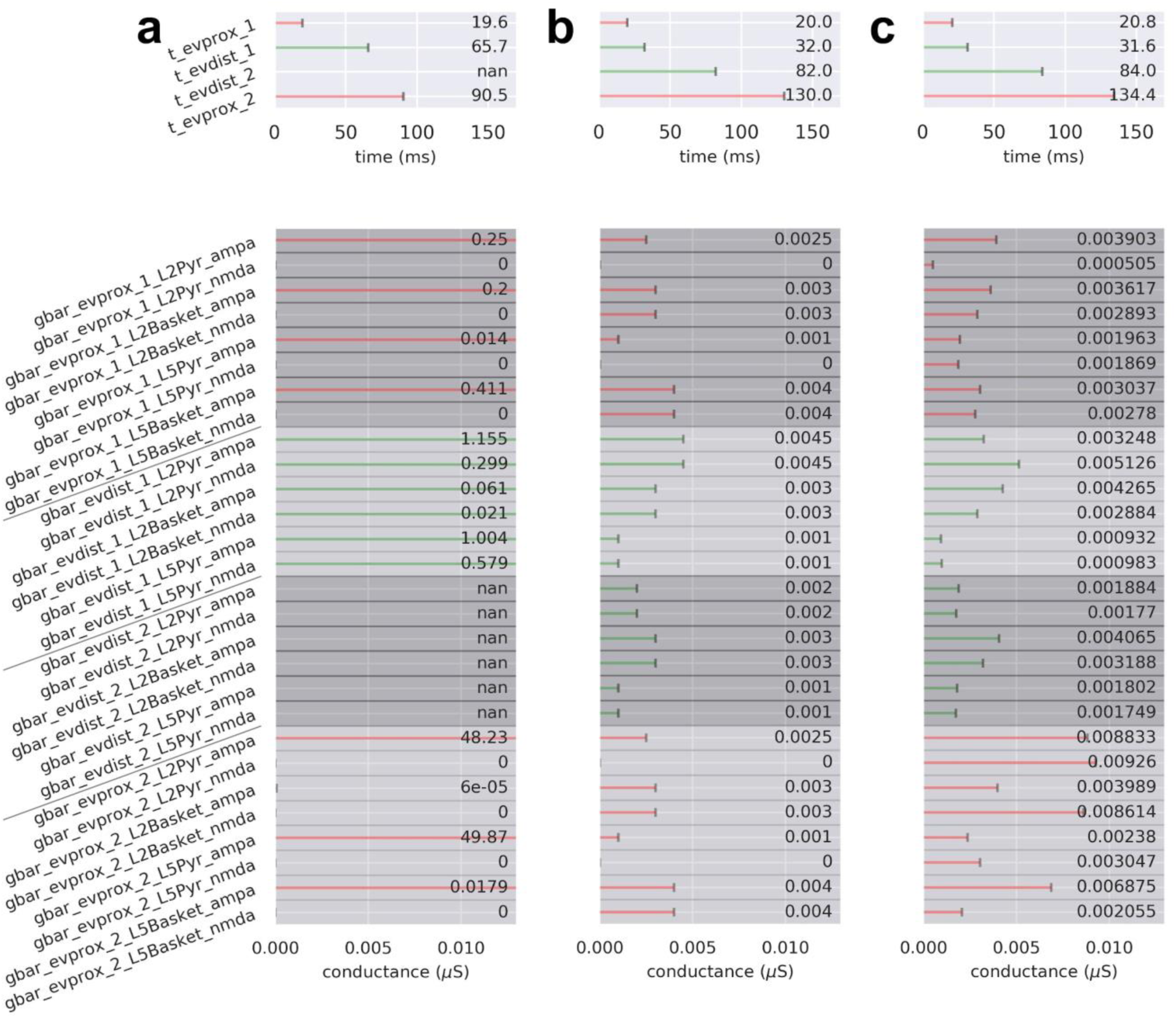
Drive parameters for the MN response simulations shown in. Figure 4. For panels (a-c), there are four possible sequential drives: an early proximal drive (evprox_1), a subsequent distal drive (evdist_1), a late distal drive (evdist_2), and late evoked proximal drive (evprox_2). Each drive has a mean spike time (prefix ‘t’; top) and synaptic weights (prefix ‘gbar’; bottom) associated with the drive’s connectivity and available neurotransmitter receptors. Parameter values denoted by ‘not a number’ (nan) correspond to an omitted drive. (a) Drive sequence consists of three drives evprox_1, evdist_1, and evprox_2 at times t=19.6±2.5 (mean±SD), 65.7±3.8, and 90.5±10.4 ms, respectively. (b) Drive sequence consists of all four possible drives: a proximal, 2 distal, and a proximal at times t=20.0±3.0, 32.0±3.0, 82.0±3.0, and 130.0±3.0 ms, respectively. (c) Post-OPS of the drive sequence in (b) with drive times t=20.8±4.1, 31.6±2.7, 84.0±4.5, and 134.4±4.4 ms, respectively.

To account for the longer duration of the P2/prolonged depression and the less pronounced return to baseline, the parameters (namely, timing and conductances) of the distal drive at ∼65 ms and subsequent proximal drive at ∼90 ms applied in the TE response simulation were hand-tuned from their initial values to values that produced a visibly close reputation to the to the MN P2/prolonged depression and rebound (Figure 4b; parameter values in Figure 6b). The distal drive occurred at t=82.0±3.0 ms and the re-emergent proximal at t=130.0±3.0 ms. Of note, the pattern of underlying spiking activity after ∼65 ms was dominated by inhibitory spiking in L2/3 and L5, whose primary effect was to pull current flow down the dendrites, decreasing the outward current to create the prolonged depression and smaller magnitude rebound (see also layer specific responses in Supplementary Figure 1b).

Following manual hand-tuning of parameters, we performed automated parameter optimization over all drive parameters to improve the fit between simulated and empirical data. This changed the parameters slightly to create a mean aggregate dipole with closer agreement to the empirical data. Of note, the optimized parameters generated a resurgence of pyramidal neuron spiking after ∼65 ms, which induced outward current via backpropagating action potentials to increase the later response and improved the final fit to the data (Figure 4c, RMSE 0.031 compared to 0.125 for pre-optimization; parameter values in Figure 6c). Since the drive sequence was preserved despite the individual mean drive times being allowed to vary in the optimization procedure, we can conclude that the order and relative timing of our drive sequence enables our neocortical column model to produce dynamics that reside near a local minimum in RMSE. We also note that there is a large amount of variability in the response near 50 ms across trials (grey traces, Figure 4c), where a small outward oriented deflection sometimes emerges and sometimes doesn’t. In other studies depicting this peak, it would be labeled N2 [or N45 as in (Wikström et al., 1996)]. Here, we did not label it due to its variable emergence across studies (see Figure 2a). Further discussion of the “N2” deflection and the mechanisms contributing to its variability are in the Discussion.

In order to understand how changes in parameters around the optimized values contribute to variance in the current dipole waveform, we performed a parameter sensitivity analysis on a key subset of the evoked proximal and distal drive parameters (evprox_1, evprox_2, evdist_1, and evdist_2), based on prior work showing these parameters created the most variance in the TE response (see Methods). Parameters of the proximal drives included their mean drive times and the maximal postsynaptic conductances of the NMDA currents only, specifically L2_pyramidal_nmda, L5_basket_nmda, and L5_pyramidal_nmda. Parameters of the distal drives included their mean drive times and the maximal postsynaptic conductances of both NMDA and AMPA currents, specifically L2_basket_ampa, L2_pyramidal_nmda, and L5_pyramidal_nmda. Each of these parameters was varied over a fixed uniformly sampled range, while keeping all other parameters fixed, and plotted the resulting net SD of the simulated current dipole contributed by each parameter (grouped by drive) at each time point (Figure 4d top left). Figure 4d (top right) shows the relative normalized proportion of SD contributed by each parameter, and (bottom) shows example waveforms for 10 different values of the single parameter that contributed the most variance over time for each drive. The early portion of the waveform (N1/P1) is controlled by the evprox_1 and evdist_1 drive parameters, as would be expected. These drives also contributed a large amount of variance during the P2/prolonged depression period but have little influence on later parts of the waveform. As would be expected based on its timing, the evdist_2 parameters controlled the amplitude of the P2 deflection and influenced the later portions of the waveform, while the evprox_2 parameters only contributed to the variance at the end of the signal. Overall, these results suggested that the earliest drives to the SI network (evprox_1 and evdist_1) have the most prominent influence on signal variance over the entire waveform.

### 3.4. HNN predicts the LE response is generated from a specific sequence of exogenous drive, such that the N1 is created from a burst of repetitive ∼40 Hz distal drives near 170 ms that induces prolonged inhibition

The LE response differs from the MN response in several ways (Figure 2). Most notable, the first deflection of the LE response occurs at ∼170 ms (LE N1), lasts ∼150 ms, and is oriented downward, reflecting inward current flow down the pyramidal neuron dendrites. Based on our prior simulation studies showing that inward currents (negative deflections) can be generated by distal drive, we hypothesized that the LE N1 could be created by a distal drive (Figure 5b). We further hypothesized that the long lasting effects of the LE N1 response could occur either through a single drive that produces a long-duration response (Figure 7a), or through multiple, repetitive drives that produce successive responses (Figure 7b). As detailed below, exploration of these alternative hypotheses suggested that a burst of four repetitive distal drives with an inter-drive-interval (IDI) of 25 ms (i.e., 40 Hz) produces the prominent, prolonged downward LE N1 deflection.

**Figure 7.**
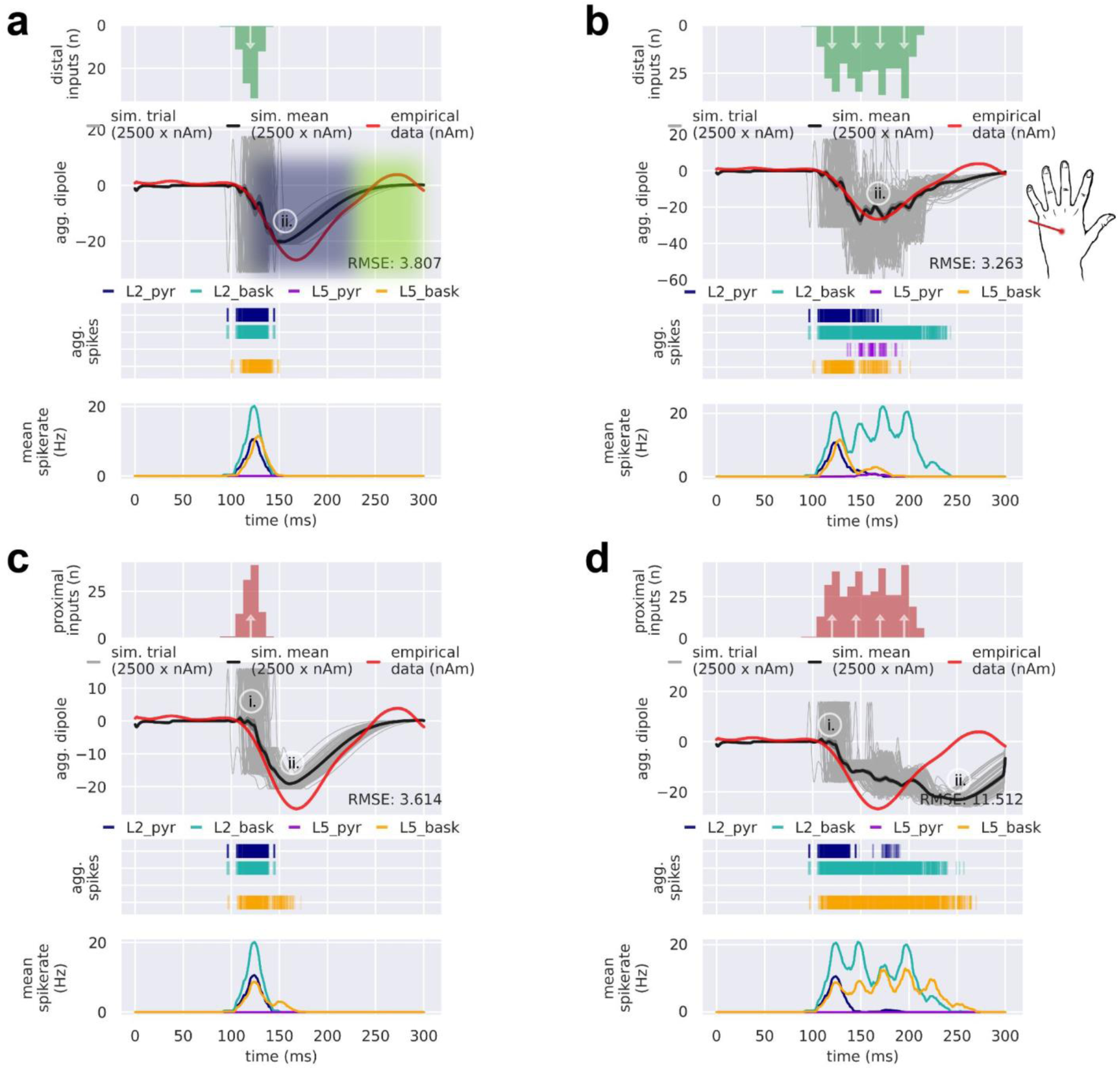
Prolonged downward current flow of the LE N1 requires strong activation of inhibitory cells. Multiple drive configurations can be employed to accomplish inhibitory basket cell activation, yet repetitive distal drives evoke a current dipole that best matches the depth and breadth of the empirical LE N1. From top to bottom, panels (a-d) each contain the input drive histogram across all 100 trials (proximal, red; distal, green), aggregate simulated dipole (individual trials, grey traces n=100; trial-mean, black trace) alongside the empirical LE response (red trace), aggregate spike raster for all neurons over all trials according to type (L2 pyramidal, navy; L2 basket, cyan; L5 pyramidal, violet; L5 basket, orange), and population-mean spikerate (i.e., the average spikerate of a neuron of a given type). (a) A single distal drive at t=120±8.0 ms ms results in a negative mean aggregate current dipole that lasts about 125 ms beginning at the onset of the LE N1. Initial excitation at the distal apical dendrite of a pyramidal cell drives current down the dendrite towards the soma. The subsequent sustained downward deflection in each simulation trial results from inhibition at the soma drawing current down the apical dendrite for a prolonged period of time. (b) A burst of four repetitive distal drives starting at t=120±8.0 ms and ending at t=195.0±8.0 ms creates a larger and more prolonged mean aggregate dipole deflection compared to the simulation in (a). (c) A single proximal drive can produce a mean aggregate current dipole similar to that of a single distal drive, except through initial excitatory input that targets pyramidal and basket cell somas. As in (a), subsequent activation of basket cells creates strong inhibition that results in a dominant somatic hyperpolarizing current. (d) A burst of four repetitive proximal drives with the same temporal profile as in (b) creates a mean aggregate dipole that extends well past 250 ms. Simulated current dipole trials in (a-d) are smoothed by convolution with a 5 ms-wide Hamming window.

Figure 7a shows a single simulated distal drive at t=120±8.0 ms that produces a long-duration response in the network. Here, the timing and postsynaptic conductances were hand-tuned starting with postsynaptic conductance values from the MN response to get an approximate fit to the LE N1 (parameters in Figure 8a). Excitation at the distal portion of the pyramidal apical dendrite initially pushed depolarizing current down towards the soma. Induced spiking activity in L2/3 and L5 inhibitory cells produced sustained hyperpolarizing downward/inward current flow via both fast and slow somatic inhibition. Note that the delayed L5 inhibitory neuron activity was induced via the excitatory connection from the L2/3 pyramidal neuron to L5 inhibitory neuron (see Figure 1 and Table 1) and the response is dominated by the activity in the longer L5 pyramidal neurons (see layer specific responses in Supplementary Figure 2a). While a single short duration distal drive produced a visably close fit to the empirical LE N1 deflection, the response did not reach the full depth and breadth of the deflection. Automated optimization of the distal drive parameters did not improve the fit and the deflection instead plateaued (see Supplementary Figure 3a).

**Figure 8.**
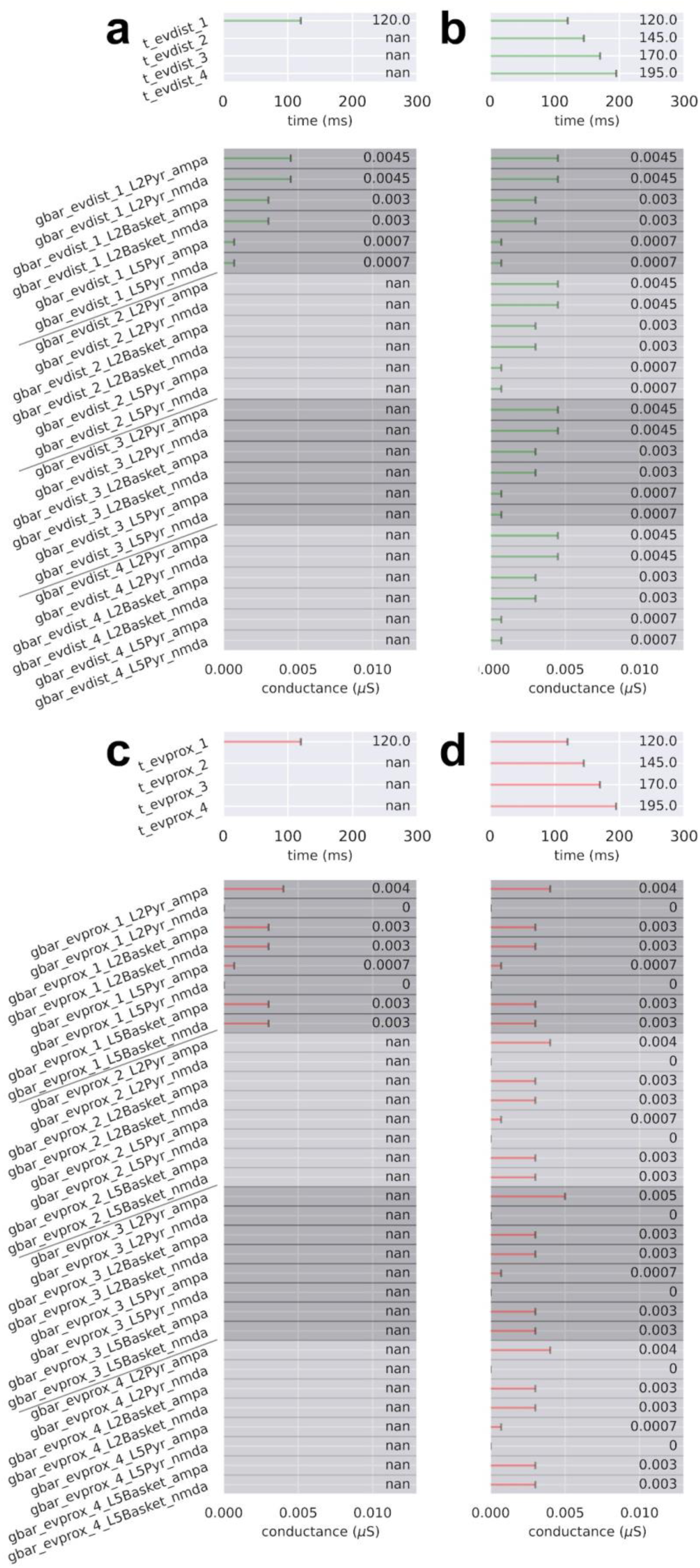
Drive parameters for the LE response simulations shown in. Figure 7. For panels (a-d), there are either one or four drives per simulation. Each drive has a mean spike time (prefix ‘t’; top) and synaptic weights (prefix ‘gbar’; bottom) associated with the drive’s connectivity and available neurotransmitter receptors. Parameter values denoted by ‘not a number’ (nan) correspond to an omitted drive. (a) Drive sequence consists of a distal drive (evdist_1) at time t=120.0±8.0 ms (mean±SD). (b) Drive sequence consists of four distal drives (evdist_1-evdist_4) at times t=120.0±8.0, 145.0±8.0, 170.0±8.0, and 195.0±8.0 ms, respectively. (c,d) Same as in (a) and (b), respectively, except with proximal instead of distal drive (evprox_1-evprox_4).

We next tested the hypothesis that a burst of repetitive distal drives could induce the full depth and duration of the LE N1 deflection (Figure 7b). Through exploration of different burst patterns (data not shown), we found that repetitive distal drives applied at regular intervals of ∼25 ms over the course of the N1 deflection produced the best fit to the empirical data. Figure 7b shows that repetitive distal drives comprised of inputs at t=120.0±8.0 (mean±SD), 145.0±8.0, 170.0±8.0, and 195.0±8.0 ms created a deeper and more prolonged mean aggregate dipole deflection (parameters shown in Figure 8b). Similar to the single distal drive, the first drive of the burst created distal depolarization of the pyramidal neurons followed by somatic hyperpolarization that together produced a downward current, With each successive distal drive, inhibitory neurons in L2/3 were re-activated creating prolonged and cumulative somatic hyperpolarization of the L2/3 pyramidal neurons that dominates the later portion of the response (see spike raster and aggregate spikerates in Figure 7b, bottom panels, and layer specific responses in Supplementary Figure 2b).

The ∼25 ms IDI of the distal drive is equivalent to ∼40 Hz gamma frequency drive. Further investigation of spiking activity in the network suggests that this drive frequency produced an optimal balance of inhibition and excitation in the L2/3 network by engaging prolonged somatic inhibition and minimizing pyramidal neuron firing (Supplementary Figure 4). The spike raster in Figure 7b shows that the population of L2/3 inhibitory neurons spikes rhythmically at ∼40 Hz. This gamma resonance is due to strong long inhibitory-to-inhibitory GABA_A_ergic connections (Table 1) that have a ∼25 ms time constant of decay, coupled with the gamma frequency distal drive to the inhibitory neurons. Gamma resonance due to fast reciprocal inhibition is an established mechanism known as Interneuron-Gamma (ING) (Tiesinga and Sejnowski, 2009; Wang and Buzsáki, 1996). This prolonged L2/3 inhibition suppresses pyramidal neuron firing in both L2/3 and L5, enabling the somatic hyperpolarization to pull current flow down the pyramidal neuron dendrites and produce a negative (inward) current dipole response. Supplementary Figure 4 shows that the ∼25 ms IDI minimized pyramidal neuron firing more effectively than several other IDI configurations (and their associated frequencies) of the repetitive drive sequence.

The predominant mechanism creating the LE N1 deflection in Figure 7a,b is prolonged pyramidal neuron somatic inhibition mediated by a single or repetitive distal drive(s). However, an open question is if a strong proximal drive to the L2/3 and L5 inhibitory neurons, absent of a distal drive, might suffice to generate the N1. We tested this alternative hypothesis by delivering a single proximal drive at t=120±8.0 ms (Figure 7c). This drive configuration produced a mean aggregate current dipole and spiking pattern similar to that of the single distal drive in Figure 7a. As with a single short duration distal input, the depth and breadth of the simulated N1 deflection did not match the empirical data. An attempt to manually deepen and prolong the N1 deflection with a burst of repetitive proximal drives, similar to the distal burst pattern in Figure 7b, did not improve fitting of the model to the experimental data and instead extended the response well past the end of the empirical LE N1 deflection (>250 ms; Figure 7d). Parameter optimization also did not adequately improve the fit with either the single or repetitive proximal drives (Figure 7c,d; Supplementary Figure 3c,d). We thus concluded that a burst of repetitive distal drives with a 25 ms IDI provided the best fit to the LE N1 response.

Given this mechanism for the LE N1 response, we next examined mechanisms that could account for subsequent rebound activity at ∼250 ms. Based on our prior knowledge of generators of outward currents (Figure 5b), we hypothesized that the rebound is produced by proximal drive. Building from the simulation in Figure 7b, Figure 9a,b shows the LE response with an additional late proximal drive at t=260±12.0 ms, before and after parameter optimization, respectively (parameter values in Figure 10a,b). The late proximal drive can generate the positive-deflecting rebound response through proximal depolarization and backpropagation of somatic spikes in the L2/3 and L5 pyramidal neurons, which together push current flow up the pyramidal neuron dendrites (post-optimization Figure 9b, middle and bottom plots). Parameter optimization preserved the temporal profile of the burst of distal drives (i.e., with a mean 25 ms IDI) and further decreased the RMSE.

**Figure 9.**
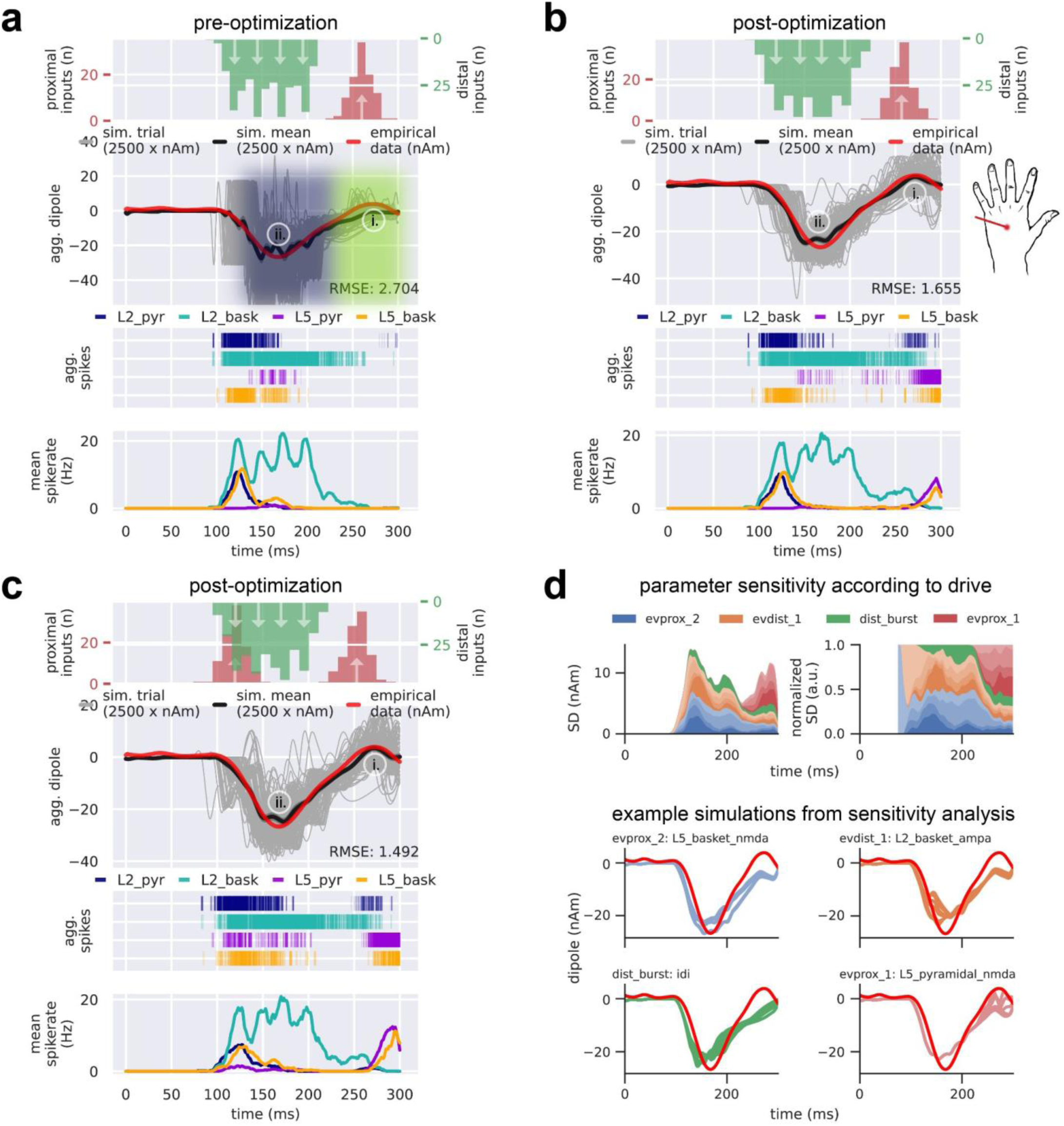
The LE N1 deflection along with its subsequent rebound can be simulated with a burst of repetitive distal drives, a late proximal drive, and an optional early proximal drive. From top to bottom, panels (a-d) each contain the input drive histogram across all 100 trials (proximal, red; distal, green), aggregate simulated dipole (individual trials, grey traces n=100; trial-mean, black trace) alongside the empirical MN response (red trace), aggregate spike raster for all neurons over all trials according to type (L2 pyramidal, navy; L2 basket, cyan; L5 pyramidal, violet; L5 basket, orange), and population-mean spikerate (i.e., the average spikerate of a neuron of a given type). (a) A manually-tuned burst of repetitive distal drives with a late proximal drive at t=260±12.0 ms simulates the N1 deflection with its subsequent rebound. (b) Parameter optimization of the drive configuration shown in (a) conserves the temporal profile of driving inputs while improving the simulation fit to empirical data. (c) Same as in (b), except optimized with an additional early proximal drive at t=120±12.0 ms. (d) Sensitivity analysis of selected parameters belonging to each of the drives illustrates the prominent and differential role respective drives have in controlling the variance of the model’s current dipole response near each major deflection. Top left: net standard deviation (SD) of the simulated current dipole contributed by each parameter at each time point. Top right: normalized net SD at each timepoint. Bottom: example simulations from the sensitivity analysis where the selected parameter was allowed to vary while keeping all others fixed. Parameters include L2_basket_ampa, L2_pyramidal_ampa, L5_basket_nmda, and mean drive time for the early proximal drive (evprox_2), L2_basket_ampa, L2_pyramidal_ampa, L2_basket_nmda, and mean drive time for the first distal drive of the distal burst (evdist_1), inter-drive-interval of mean drive times for the distal burst (dist_burst), and L2_pyramidal_amp, L5_pyramidal_amp, L5_pyramidal_nmda, and mean drive time for the late proximal drive (evprox_1). Simulated current dipole trials in (a) are smoothed by convolution with a 5 ms-wide Hamming window and trials in (b-d) are smoothed by convolution with a 20 ms Hamming window.

**Figure 10.**
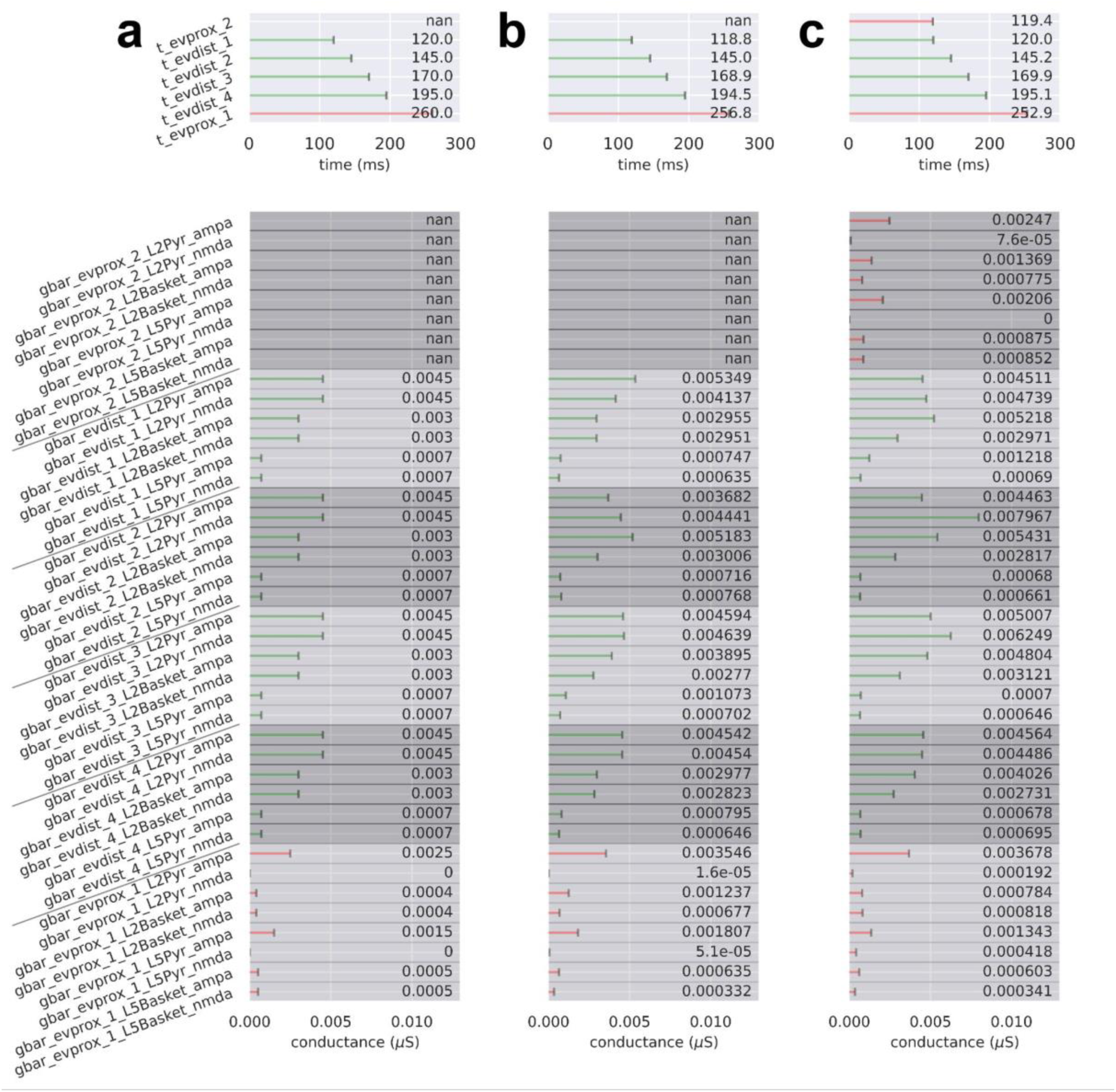
Drive parameters for the LE response simulations shown in. Figure 8. For panels (a-c), there are six possible sequential drives: four distal drives (evdist_1-evdist_4), a proximal drive responsible for producing the rebound (evprox_1), and an optional early proximal drive (evprox_2). Each drive has a mean spike time (prefix ‘t’; top) and synaptic weights (prefix ‘gbar’; bottom) associated with the drive’s connectivity and available neurotransmitter receptors. Parameter values denoted by ‘not a number’ (nan) correspond to an omitted drive. (a) Drive sequence consists of distal drives evdist_1-evprox_4 and evprox_1 at times t=120.0±8.0 (mean±SD), 145.0±8.0, 170.0±8.0, 195.0±8.0, and 260.0±12.0 ms, respectively. (b) Post-OPS of the drive sequence in (a) with drive times t=118.8±10.0, 145.0±10.0, 168.9±9.2, 194.5±11.5, and 256.8±12.2 ms, respectively. (c) Post-OPS of the drive sequence in (a) except with an additional early proximal drive, exprox_2, and drive times t=119.4±12.0, 120.0±10.0, 145.2±10.0, 169.9±8.8, 195.1±10.1 and 252.9±12.3 ms, respectively.

The sequence of distal followed by proximal drive that generated the LE response in Figure 9a,b is remarkably different than the established cortical response to innocuous somatosensory stimulation, which begins with a feedforward drive via lemniscal thalamic input (e.g., as with the MN N1 response in Figure 4). However, as a single proximal drive was able to replicate some features of the LE N1 response (Figure 7c) and the in vivo laminar profile of the LE response is unknown, we also tested if a proximal-first sequence, similar to the MN drive sequence, could in fact produce the LE response. We tested this by adding an early proximal drive at t=120.0±12.0 ms to the existing drive sequence for the LE response from Figure 9a and then running parameter optimization (Figure 9c; parameters in Figure 10c). The net result of adding the early proximal input was an increase in early L5 pyramidal spiking at ∼125 ms and a slightly improved fit to the empirical waveform post-optimization. We concluded that this early proximal input is possible; however, its necessity is unknown. As addressed further in the Discussion, the model results provide several targeted predictions on the multiscale cortical dynamics underlying these alternative mechanisms that can be tested further with invasive recordings or other imaging modalities (e.g., early firing in the L5 pyramidal neurons would be consistent with an early proximal input).

As a final step, we performed a parameter sensitivity analysis to examine how changes in parameters around the final optimized values contributed to variance in the current dipole waveform (Figure 9d). Once again, we focussed on a key subset of the evoked proximal and distal drive parameters (see Methods). Parameters of the early evoked proximal drive (labeled evprox_2, since it was the second drive added), included the mean drive time and the maximal postsynaptic conductance of AMPA and NMDA, specifically L2_basket_ampa, L2_pyramidal_ampa, and L5_basket_nmda. Parameters of the late evoked proximal drive (evprox_1) included the mean drive time and AMPA and NMDA currents onto the pyramidal neurons, specifically L2_pyramidal_amp, L5_pyramidal_amp, and L5_pyramidal_nmda. Parameters of the distal drives included the mean drive times of the burst of inputs and their average IDI, as well as the maximal postsynaptic conductance of AMPA and NMDA currents, specifically L2_basket_ampa, L2_pyramidal_ampa, and L2_basket_nmda. Figure 9d shows the resulting net SD of the simulated current dipole contributed by each parameter (grouped by drive) at each time point (top left), the relative normalized proportion of SD contributed by each parameter (top right), and example waveforms for 10 different values of the single parameter that contributed the most variance over time for each drive (bottom). As expected, variance in the early portion of the waveform (N1) is controlled by the earliest drives (evprox_2 and evdist_1). Variance from these drives also persisted into the rebound period, while the later proximal input (evprox_1) influenced only the rebound period. Overall, these results suggested once again that variance in each peak is dominated by the parameters of the drive that is closest to it, and that the earliest drives have the most prominent influence over the entire waveform.

### 3.5. The simulated LE response exhibits robust gamma activity during the N1 deflection

Recent studies have shown that the early response to noxious laser stimulation includes robust ∼40 Hz and ∼80 Hz gamma activity in human EEG sensors near SI (Heid et al., 2020; Hu and Iannetti, 2019), the latter of which is also prominent in rodent ECoG and intracortical microelectrode recordings (Hu and Iannetti, 2019; Yue et al., 2020). Given that inhibitory cell firing at ∼40 Hz helps create the negative LE N1 deflection in the model simulations (Figure 7b, Figure 9, and Supplementary Figure 4), we tested if the frequency response of the unsmoothed simulated current dipole near the LE N1 deflection exhibits 40 Hz activity, thus providing a potential circuit explanation for the event-related synchronizations observed in prior empirical studies. Indeed, prominent ∼40 Hz spectral activity was present in single-trial TFRs (Figure 11a-c) and the trial mean TFR (n=100 trials averaged in the spectral domain; Figure 11d) of the simulated LE response in Figure 9c, along with peaks at ≤5 Hz (theta-band) and ∼80 Hz (high gamma-band). The 80 Hz activity emerges from the time course of the sharp deflections in the waveform that each last at most ∼12.5 ms. The 40 Hz activity reflects the periodic nature of the downward (inward) dipole deflections occurring at ∼25 ms intervals from somatic inhibition, as described above, and the ≤5 Hz activity comes from the slow downward component of the LE N1 deflection that lasts ∼100 ms. These results were consistent across the simulations with and without an early proximal drive as presented in Figure 9b,c (comparison not shown).

**Figure 11.**
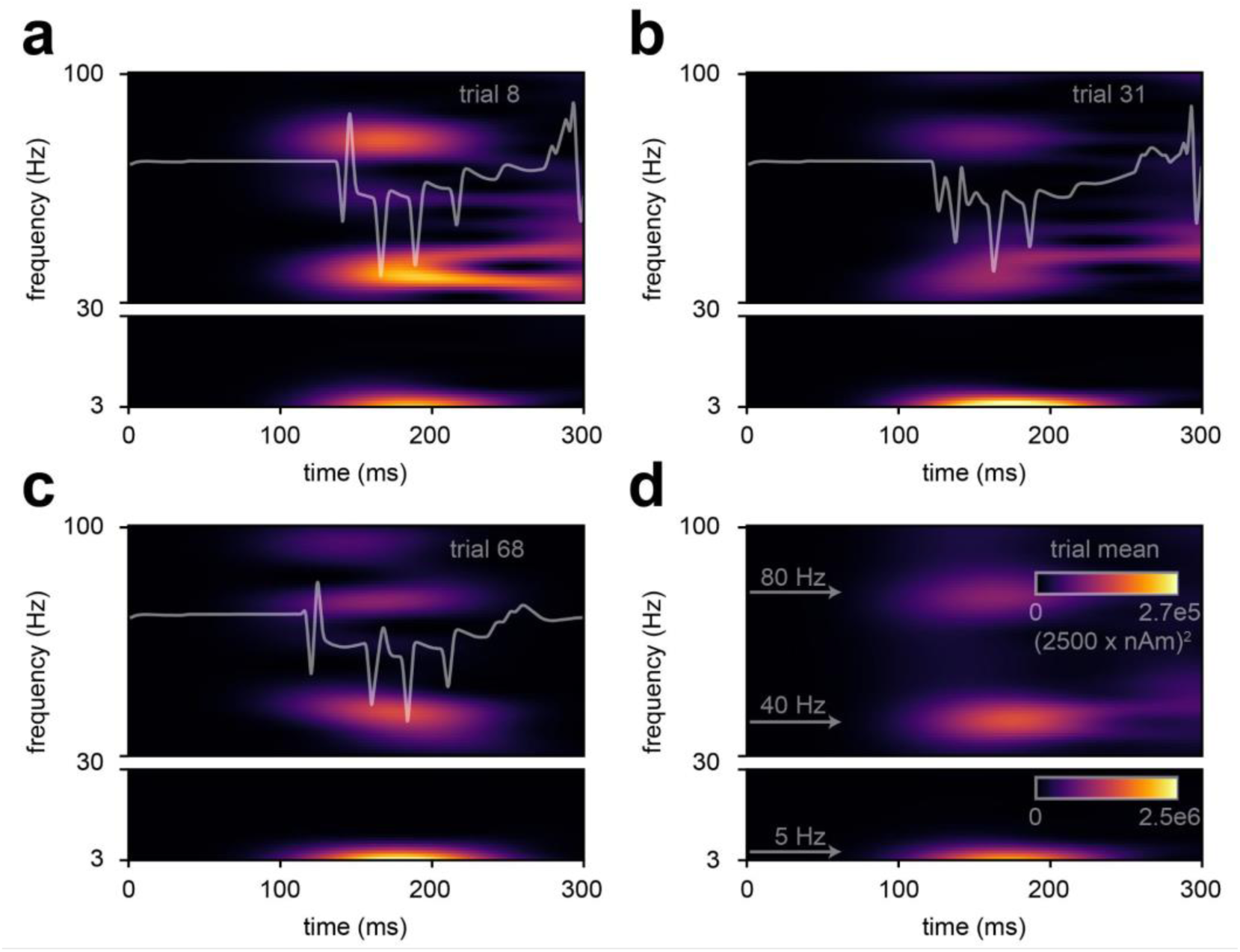
Repetitive distal drives in the optimized LE N1 simulation produce 40 Hz resonant spectral activity in the model SI circuit. (a-c) Time-frequency response (TFR) of three representative simulation trials. (d) Trial-mean TFR of individual simulation trials (n=100) reveals LE event related spectral activity at ∼5, ∼40, and ∼80 Hz. Note that the TFR was calculated from simulation trials with minimal smoothing (i.e., by convolution with a 5 ms Hamming window) to preserve the high frequency components of each time course.

## 4. Discussion

M/EEG evoked responses from median nerve and laser stimulation are widely studied to gain insight into non-painful and painful somatosensory processing. The majority of prior studies describe how evoked response features such as peak timing and amplitudes vary with stimulation conditions and/or perception. Here, we address the open question of the detailed neocortical circuit origin of the deflections in innocuous MN and noxious LE responses found in macroscale neural recordings. Analyzing data from prior published studies, we found that the peak timings and orientations of source-localized responses in SI were conserved across studies and showed that each peak can be interpreted in terms of the corresponding current flow up and down pyramidal neuron apical dendrites (i.e., out of or into the cortex). We used the HNN simulation software, whose foundation is a laminar model of the neocortex under thalamocortical and cortico-cortical drive, to predict the detailed cell and circuit mechanisms underlying this current flow.

Building on prior HNN studies of the mechanisms of generation of source-localized evoked responses, particularly those from somatosensory cortex (Jones et al., 2009, 2007; Kohl et al., 2021; Law et al., 2021; Sliva et al., 2018; Ziegler et al., 2010), we tested and refined hypotheses on the multiscale generation of previously recorded MN and LE responses. As summarized in Figure 12, HNN modeling predicted that the mechanisms of the MN response are similar to those of the tactile evoked response and generated from a sequence of layer specific feedforward (i.e., proximal drive from lemniscal thalamus), feedback (i.e., distal drive from non-lemniscal thalamus or higher order cortex) and re-emergent feedforward drive to the local network, with the required addition of early distal drive at ∼30 ms to generate the MN P1 (P30) peak. Unlike the MN response, the first peak in the LE response (N1) represents an inward current, which modeling predicted emerges from an excitatory synaptic input to the supragranular layers (i.e., distal). Additionally, this distal input consisted of repetitive drives at 40 Hz that generated sustained inhibition to create the downward deflection and produced gamma frequency activity in the single trial simulations in the model.

**Figure 12.**
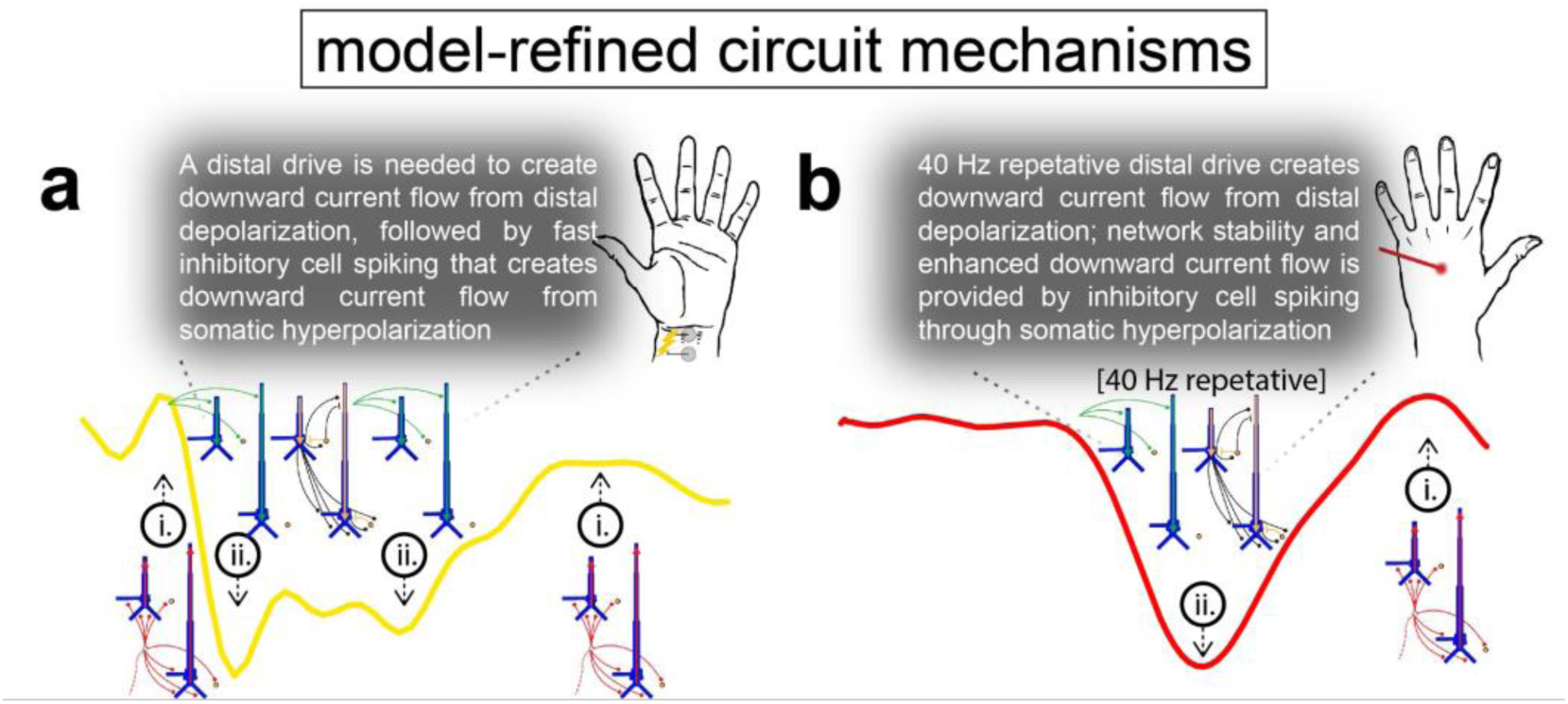
Final model-refined circuit mechanisms that underlie each deflection of the respective MN and LE responses.

Overall, the HNN model’s ability to produce both the MN and LE responses by only modulating the sequence of exogenous drives to the local network, with all local network parameters held constant, supports the notion that the canonical features of a laminar neocortical network are capable of producing both innocuous and noxious responses. It should be noted that we make no claims about the relative size, number or overlap of the neuron populations responsible for generating the MN and LE responses within SI; rather we only posit that both evoked responses can emerge from canonical features of the SI network. We also make no specific claims about mechanisms underlying perception but focus on the mechanisms that produce features of the MN and LE responses, such as peak timings and orientations, that are conserved across studies (Figure 2).

Below, we place the multiscale model-derived predictions within the context of prior studies investigating the neural mechanisms of MN and LE responses and the known anatomy and physiology of thalamocortical and cortico-cortical circuitry. We emphasize predicted circuit dynamics that can guide further testing with invasive recordings or other imaging modalities.

### 4.1. Model predictions extend prior understanding of the circuit mechanisms of the MN response

Consistent with previous studies, the modeling results here demonstrate that the first deflection of the MN response can be produced by feedforward excitatory synaptic drive from thalamus to the granular layers of SI at ∼20 ms. In the model, this input propagates directly to the proximal dendrites of pyramidal neurons in supra- and infragranular layers to induce current flow up the pyramidal apical dendrites (Cauller and Kulics, 1991; Jones et al., 2007; Lipton et al., 2006; Peterson et al., 1995; Schroeder et al., 1995; Wikström et al., 1996). This drive likely originates in the ventrobasal thalamus (Cauller and Kulics, 1991; Jones et al., 1979; Yamada et al., 1984) as part of the lemniscal pathway that receives activation from fast Aβ fibers in the peripheral nerves (Burke et al., 1975).

Following the initial MN N1 (N20) deflection, our model predicts that an early distal drive at ∼30 ms is necessary to produce the downward MN P1 (P30). This novel prediction deviates from an alternative notion in the literature that assumes the human M/EEG P1 deflection from median nerve (Wikström et al., 1996) and tactile stimulation (Jones et al., 2007), also known as the downward-pointing EEG/local field potential (LFP) N1 in rats (Bruyns-Haylett et al., 2017), originates from somatic hyperpolarization of pyramidal neurons via proximal “feedforward” inhibition. While the precise pre-synaptic source of this predicted ∼30 ms distal drive is unknown, likely candidates include the ventromedial thalamus and/or the secondary somatosensory cortex (SII) given that they both process somatosensory signals and project into the superficial layers of SI (Cauller and Kulics, 1991; Friedman, 1983). Thus, we further predict that either the innocuous MN signal from the periphery branches somewhere in its ascending pathway, or there is a rapid thalamocortical loop of activity that creates both proximal and distal drive within the first ∼30 ms post-stimulus.

In addition to creating the MN N1 and P1 deflections, the precise timing of the early proximal and distal drives relative to each other created an optimally excitable state that allowed backpropagation of pyramidal neuron spikes to generate a small outward-oriented rebound before a subsequent distal drive at ∼85 ms induced the prominent P2/prolonged depression. Historically, this small outward rebound has been referred to as the “N2”, but was not labeled here due to its variability (see Figure 2). In prior studies, the mechanistic origins of the MN N2 and subsequent P2 (sometimes also referred to as N45 and P60) have been disputed over as emerging from either excitation followed by inhibition at the pyramidal soma (Gardner et al., 1984; Wikström et al., 1996), excitation at the pyramidal distal apical dendrite via a cortico-cortical projection that then causes current to passively flow down the dendrite toward the soma (Cauller and Kulics, 1991), or a combination of the two (Peterson et al., 1995). The model predictions here are supportive of a combined mechanism, where local network spiking in the L5 pyramidal cells emerges from local excitatory synaptic interactions and creates the small outward oriented deflection (N2) while simultaneous prolonged somatic inhibition keeps this defection below zero. Finally, a later excitatory distal drive creates focal downward current flow during the P2/prolonged depression.

The model results may also help explain the large amount of variability in whether or not the N2 response is observed across studies (see Figure 2a). Individual simulation trial responses show enhanced variability between 50-100 ms (see also grey traces in the second panel of Figure 4c). This variability comes from the stochastic nature of the mean arrival time of the initial proximal and distal drives before 50 ms, which were chosen from a Gaussian distribution (see red and green histograms in Figure 4c). This stochasticity produces varying amounts of GABAergic inhibitory tone onto the population of L5 pyramidal cells based on the elapsed time between the proximal and distal drive. On trials with more inhibition, which is denoted by the brief suppression of population spiking across all cell types (Figure 4c), all L5 pyramidal neurons are suppressed and the N2 peak remains nearly flat, whereas on trials with less inhibition, L5 pyramidal neuron spiking creates backpropagating current and an upward N2 peak. The N2 peak, however, remains below zero because of the competing somatic inhibition. This prediction suggests that if MN stimulation is repeated at regular sub-second intervals, residual inhibition at t>150 ms will reduce the excitatory impact of the next stimulus’ early proximal and distal drive. Such an impairment in excitation tips the balance of network spiking in favor of reduced pyramidal neuron spiking and GABA-mediated inhibition that would normally produce large N1 and P1 deflections while minimizing the N2 rebound. Consistent with the results of prior research that observed the impact of repetitive MN stimulation, this mechanism predicts MN N1 and P2 attenuation and MN N2 amplification with shortening of the inter-stimulus interval and may point toward the use of MN N1, P1, and N2 deflections as a proxy for GABAergic tone (Wikström et al., 1996). Furthermore, hysteresis of these deflections due to repetitive stimulation might contribute to a central representation for perceived sensory adaptation (Graczyk et al., 2018).

### 4.2. Model generates novel predictions on the circuit mechanisms of the LE response, including a mechanistic understanding of evoked gamma oscillations during nociceptive processing

In contrast to the MN response, the first deflection of the LE response reflects an inward superficial-to-deep current which the model predicts emerges from repeated excitatory drives to supragranular layers that recruits sustained inhibition. These drives push current down the dendrites of the L2 pyramidal neurons and activate L2/3 and L5 soma-targeting inhibitory cells that simultaneously induce current flow down the pyramidal neuron dendrites. The exogenous source of these distal drives is currently unknown and is likely different from the ventrobasal lemniscal thalamic drive inducing the initial feedforward MN response. Not only does the LE response begin at a greater latency from stimulation onset (∼170 ms), reflecting in part that it is transmitted through slow Aδ fibers from the periphery to the spinal cord (Bromm and Treede, 1984; Treede et al., 1995, 1988), but early LE supragranular drive likely hales from non-sensory-specific ventromedial thalamus (VM) (Borszcz, 2006; Desbois and Villanueva, 2001; Glenn et al., 1982; Monconduit et al., 2003, 1999) or from the secondary somatosensory cortex (SII) (Baumgärtner et al., 2006; Iannetti et al., 2005; Ploner et al., 1999; Qiu et al., 2006; Treede et al., 2003; Vierck et al., 2013). As such, the source of this initial drive may overlap with the exogenous source of drives that produce later MN deflections (Cauller and Kulics, 1991). While our model simulations predict that exogenous drive targeting the superficial layer is the primary mechanism through which the LE N1 deflection is generated, it remains unclear whether or not a feedforward input to the granular layer may also be present (Figure 9c). This early feedforward drive could originate from the ventral posterolateral nucleus (VPL) of the thalamus (Treede et al., 2003; Vierck et al., 2013). The simulation results produce distinguishing micro-circuit features that can help disambiguate the existence of this feedforward drive, such as early spiking activity in L5 pyramidal neurons (compare Figure 9b and 9c).

Further, the model predicts that a burst of repetitive distal drives spaced at ∼25 ms intervals (i.e., low gamma period) produces the LE N1 deflection. Model simulations found that this burst of drives created non-phase-locked high (∼80 Hz) and low (∼40 Hz) gamma activity. This points toward a mechanistic link between the phase-locked LE N1 deflection and non-phase-locked gamma event-related synchronization (ERS) observed in prior studies (Heid et al., 2020). These results are also consistent with recent work that found significant spike-field coherence between single unit inhibitory neurons in the superficial layers of SI and nearby LFP/ECoG gamma-frequency ERS (Yue et al., 2020).

While our study does not make any claims about circuit mechanisms underlying perceived pain intensity with noxious LE stimulation, several of the predicted mechanisms can be related to prior pain studies. Numerous studies have observed that single-trial measurements of the SI LE response amplitude correlate with subjective pain ratings (Hu and Iannetti, 2019; Huang et al., 2013; Peng et al., 2018; Timmermann et al., 2001). However, our MN nerve results and other studies show that other forms of sensory stimuli (e.g., somatosensory, auditory, visual) can evoke similar waveforms features as the LE response, albeit in different brain regions (Mouraux and Iannetti, 2009) and that they grade with perceptual intensity (Hu and Iannetti, 2019). The implied conclusion has been that the LE response waveform lacks the ability to encode nociceptive-specific processing and that peaks of the LE response are rather indirect measures of nociceptive processing that are easily biased by saliency (Iannetti et al., 2008). Perhaps a more direct measure of pain intensity is the magnitude of induced gamma-band oscillations near the time of the LE N1. The magnitude of gamma-band ERS correlates with perceived noxious stimulus intensity (Peng et al., 2018), is independent of saliency (Zhang et al., 2012), and predicts nociceptive-specific perception both within and across subjects (Heid et al., 2020; Hu and Iannetti, 2019). Further evidence suggests that spontaneous bursts of EEG gamma-band activity emerging from EEG electrodes over SI can differentiate human participants who are experiencing chronic pain from those who are not (Levitt et al., 2020).

As mentioned above, our results provide a detailed interpretation of the circuit mechanisms creating the gamma ERS. This mechanism relies on repeated exogenous ∼40 Hz gamma-frequency activation of L2/3 inhibitory neurons. Remarkably, this prediction is consistent with the finding that 40 Hz optogenetic drive of parvalbumin-positive inhibitory interneurons (PV) in mice led to nociceptive hypersensitivity and aversive behavior to innocuous contextual cues (Tan et al., 2019). While this finding did not motivate the choice of gamma frequency drive demonstrated in the present study, the cohesion with many prior studies suggests interneuron mediated gamma oscillations are a marker of painful nociceptive processing (Tan et al., 2019; Yue et al., 2020).

## 5. Conclusions

This study applied the HNN neural modeling software to make multi-scale predictions about neural mechanisms producing human SI evoked responses to innocuous MN and noxious LE stimuli. Our modeling predictions complement prior research that detail the neural dynamics of sensory processing and provide several targets for further testing of novel circuit mechanisms with invasive electrophysiological recordings (e.g., multi-electrode array recordings or ECoG) or other imaging modalities [e.g., layer resolved MEG (Bonaiuto et al., 2021) or magnetic resonance spectroscopy]. Our results lay the foundation for future investigations using median nerve and laser stimulation to study circuit mechanisms associated with non-painful and painful percepts and may help guide novel therapeutics for pathological somatosensation such as somatic sensitivity and acute neuropathic pain.

## Acknowledgments

We thank Lauri Parkkonen for helpful discussion regarding the MEG MN response data which we conducted HNN analysis on. We also thank Cedric Lenoir and André Mouraux for providing EEG sensor-space MN and LE response data used in preliminary analysis for this study. HNN simulations were conducted using computational resources and services at the Center for Computation and Visualization, Brown University. This study was supported by the National Institute of Biomedical Imaging and Bioengineering (R01EB022889), National Institute of General Medical Sciences (P20GM103645, S10OD025181), and National Institute of Neurological Disorders and Stroke (R01NS108414), and the National Institute of Mental Health (T32MH115895) of the National Institutes of Health. The content is solely the responsibility of the authors and does not necessarily represent the official views of the National Institutes of Health.

---

**Supplementary Figure 1.**
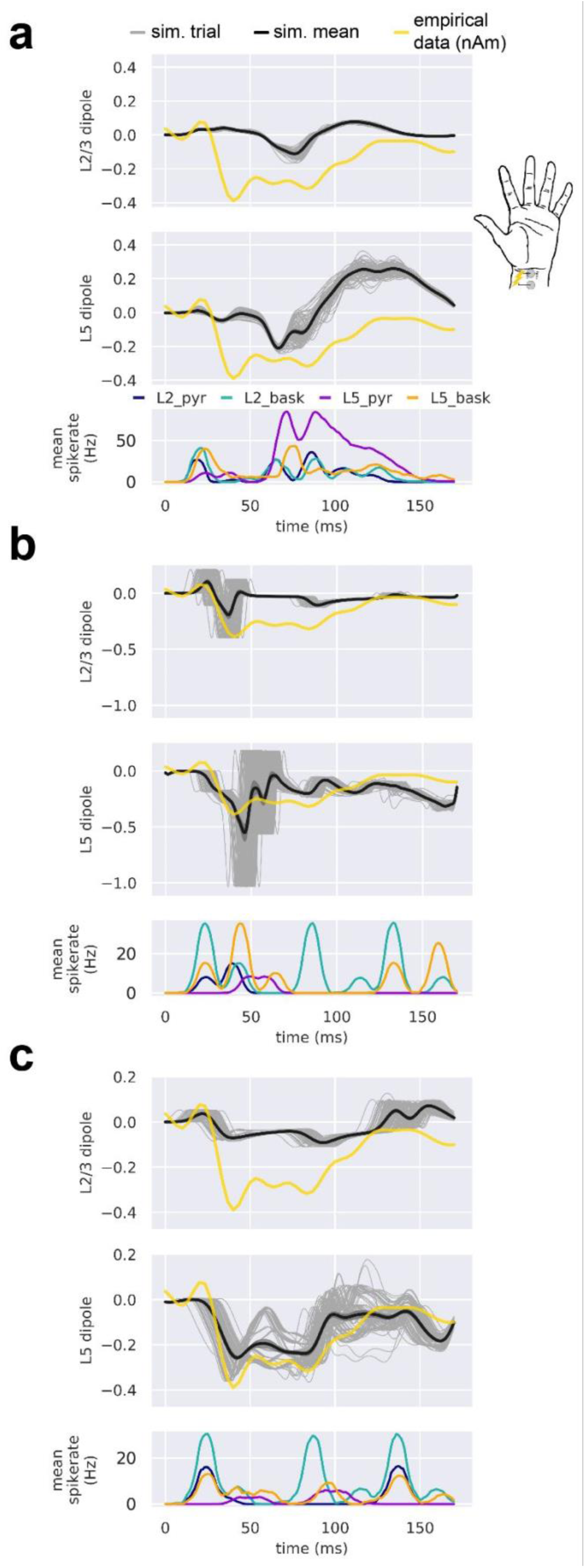
Pyramidal neurons from L2/3 and L5 contribute differently to the simulated current dipole of the MN response. From top to bottom, each panel contains the simulated dipole (individual trials, grey traces n=100; trial-mean, black trace) from L2/3 pyramidal neurons alongside the empirical MN response (yellow trace), the simulated dipole from L5 pyramidal neurons, and the population-mean spikerate (i.e., the average spikerate of a neuron of a given type). Panels (a-c) correspond, respectively, to Figure 4a-c.

**Supplementary Figure 2.**
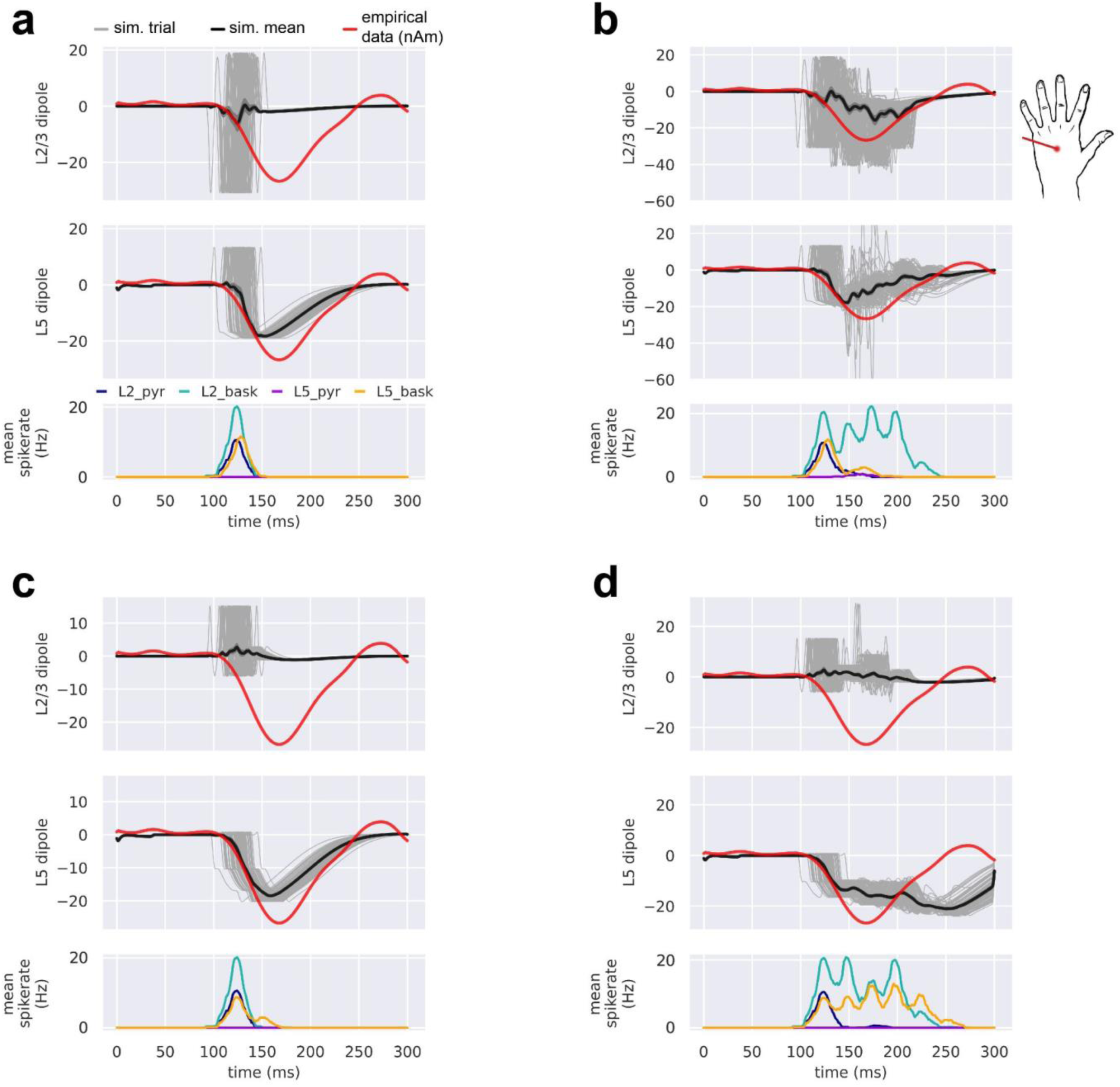
Pyramidal neurons from L2/3 and L5 contribute differently to the manually-tuned simulated current dipole of the LE response. From top to bottom, each panel contains the simulated dipole (individual trials, grey traces n=100; trial-mean, black trace) from L2/3 pyramidal neurons alongside the empirical LE response (red trace), the simulated dipole from L5 pyramidal neurons, and the population-mean spikerate (i.e., the average spikerate of a neuron of a given type). Panels (a-c) correspond, respectively, to Figure 7a-c.

**Supplementary Figure 3.**
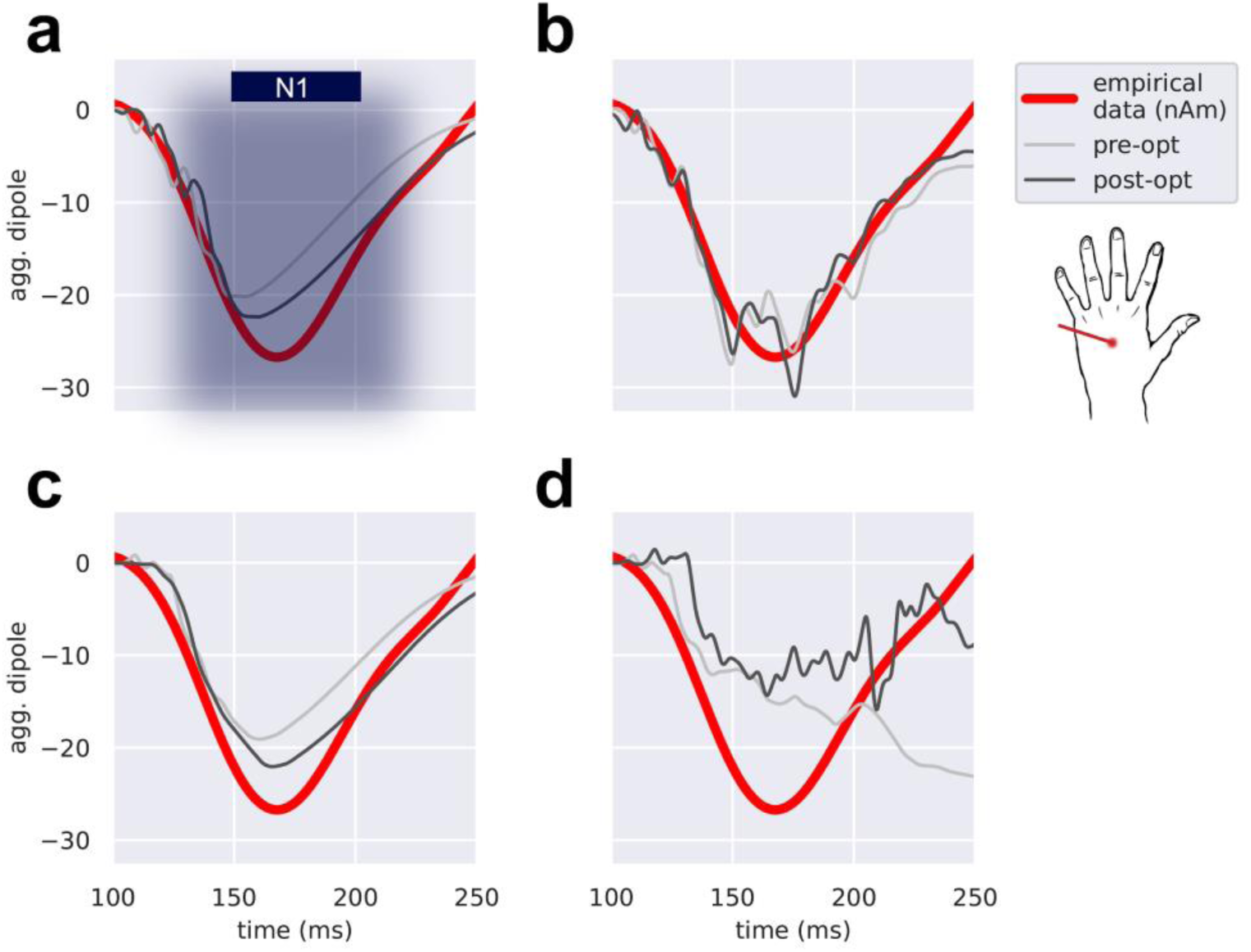
Parameter optimization provides minimal improvement to the manually-tuned drive configurations shown in Figure 7. Except for the case with repetitive distal drives (b), the trial-average aggregate current dipole before (pre-opt, grey) and after (post-opt, black) optimization for all other drive configurations including a single distal drive (a), a single proximal drive (c), and repetitive proximal drives (d) fail to simulate the depth and breadth of the empirical LE N1 deflection (red).

**Supplementary Figure 4.**
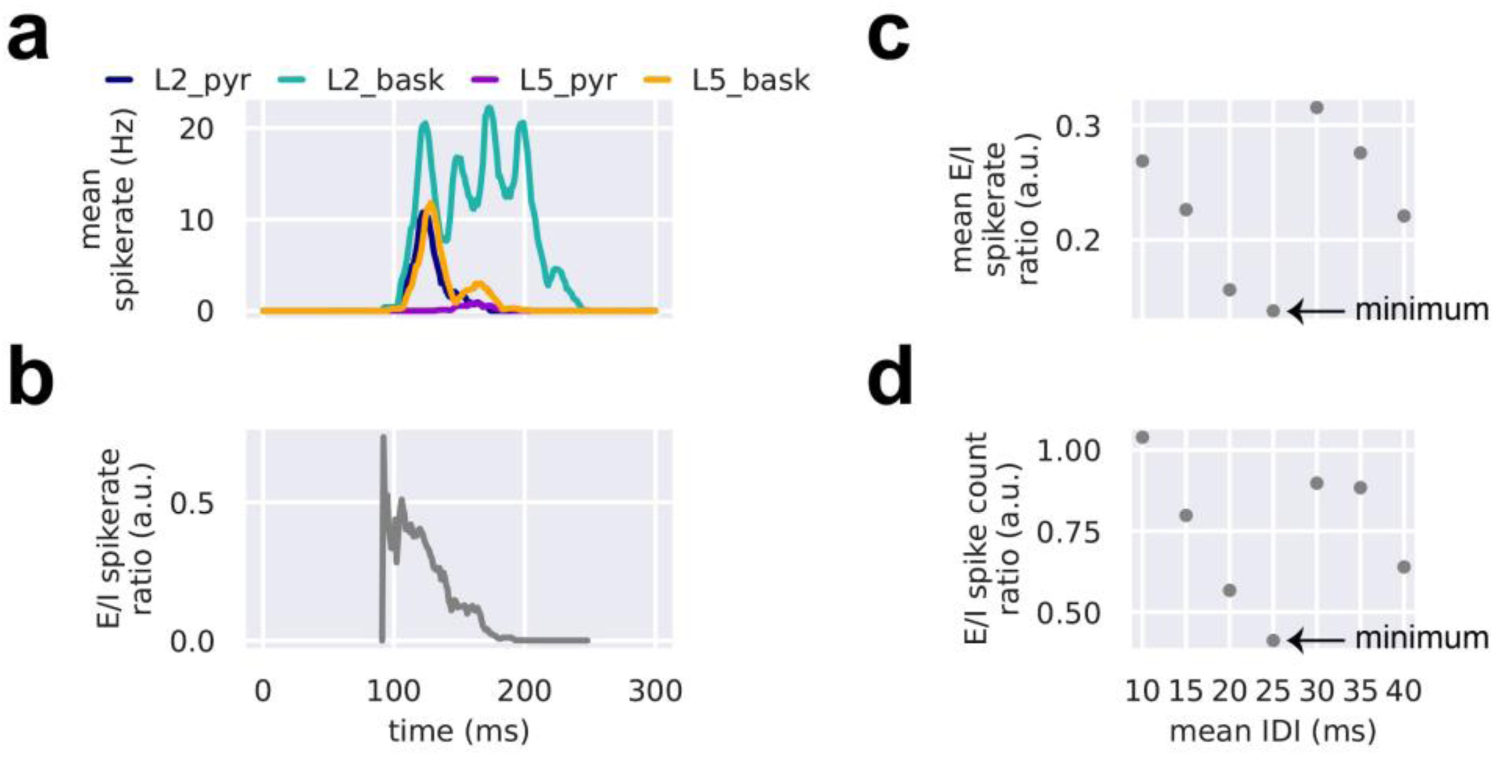
A burst of repetitive distal drives delivered at mean intervals of 25 ms minimizes network excitation relative to network inhibition. Such an excitation/inhibition (E/I) balance maximizes downward current flow in the pyramidal apical dendrites while preventing runaway excitation. (a) With a mean 25 ms inter-drive-interval (IDI), repetitive distal drives recruit large amounts of L2/3 inhibitory basket cell spiking at corresponding intervals while largely diminishing the spikerate of other (i.e., L2/3 pyramidal, L5 pyramidal, and L5 basket) cell populations. (b) Also with a mean 25 ms IDI, the ratio of the aggregate mean excitatory (L2/3 + L5 pyramidal) cell spikerate divided by inhibitory (L2/3 + L5 basket) cell spikerate is always less than 1 and attenuates with successive distal drives from 120-195 ms. (c) Across simulations with mean IDIs ranging from 10-40 ms, the time-average E/I spikerate ratio is minimal with an IDI of 25 ms. (d) Same as (c), except instead of mean E/I spikerate ratio, we show the total E/I spike count ratio across all cells, all trials, and all time of the simulation.

**Supplementary Figure 5.**
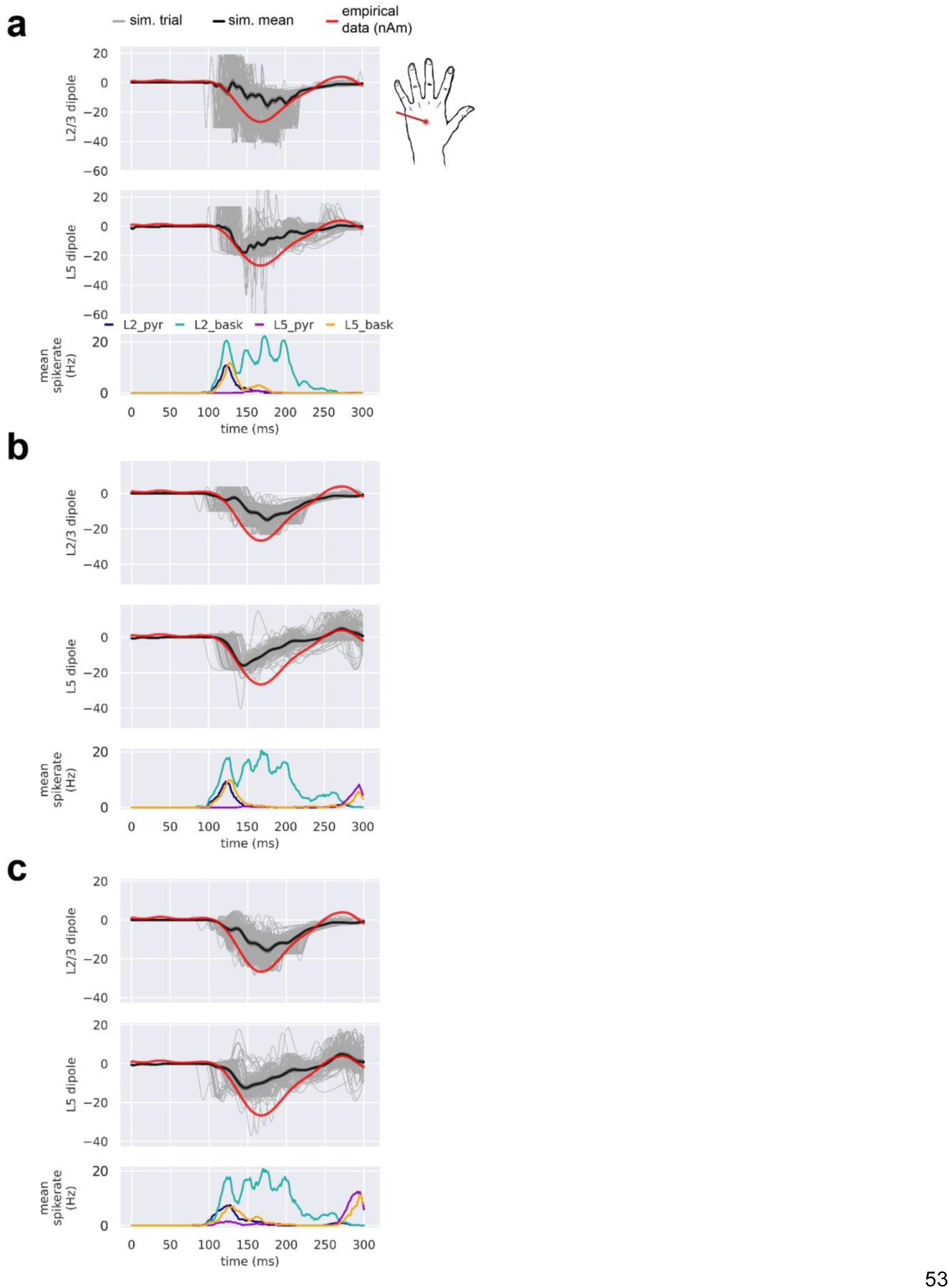
Pyramidal neurons from L2/3 and L5 contribute differently to the optimized simulated current dipole of the LE response. From top to bottom, each panel contains the simulated dipole (individual trials, grey traces n=100; trial-mean, black trace) from L2/3 pyramidal neurons alongside the empirical LE response (red trace), the simulated dipole from L5 pyramidal neurons, and the population-mean spikerate (i.e., the average spikerate of a neuron of a given type). Panels (a-c) correspond, respectively, to Figure 9a-c.

